# Transport and InsP8 activation mechanisms of the human inorganic phosphate exporter XPR1

**DOI:** 10.1101/2024.10.04.616715

**Authors:** Qinyu Zhu, Madeleine F. Yaggi, Nikolaus Jork, Henning J. Jessen, Melinda M. Diver

## Abstract

Inorganic phosphate (Pi) has essential metabolic and structural roles in living organisms. The Pi exporter, XPR1/SLC53A1, is critical for maintaining cellular Pi homeostasis. When intercellular Pi is high, cells synthesize inositol pyrophosphate (1,5-InsP_8_) – a signaling molecule that is required for XPR1 function. Inactivating mutations of XPR1 lead to brain calcifications causing neurological symptoms that include migraine, movements disorders, psychosis, and dementia. Distinct cryo-electron microscopy structures of dimeric XPR1 and functional characterization define the substrate translocation pathway and delineate how binding of InsP_8_ initiates the transport cycle. InsP_8_ binding rigidifies the intracellular SPX domains with InsP_8_ acting as a bridge between dimers and the SPX and transmembrane domains. When locked in this state, the C-terminal tail is sequestered revealing the entrance to the transport pathway, thus explaining the obligate roles of the SPX domain and InsP_8_. Together, these findings advance our understanding of XPR1 transport activity and expand opportunities for rationalizing disease mechanisms and therapeutic intervention.

## Introduction

Inorganic phosphate (Pi) is essential for all cells, serving critical roles in bioenergetics (ATP, GTP), metabolic regulation (i.e., glycolysis and oxidative phosphorylation), intracellular signaling pathways, cell proliferation (as part of DNA and RNA), and formation of structures such as bones and membranes^1^. Pi is also one of the most pervasive inhibitors of cellular enzymes, such as phosphatases where it competes for binding to the active site. As one would expect for a metabolite of such central importance, the intracellular concentration and total body content of Pi are highly regulated^2^. Transporters that facilitate Pi translocation across cellular membranes play a crucial role in this regulation. In humans, there are two protein families of sodium (Na^+^)-dependent Pi importers, SLC20 and SLC34^3^. A single Pi exporter has been identified – XPR1 (Xenotropic and Polytropic Retrovirus Receptor 1), also known as SLC53A1, as it belongs to the solute carrier (SLC) family of membrane transporter proteins^4^.

In eukaryotes, including yeast, plants, and animals, control of Pi homeostasis is, in part, conferred through a family of SPX domain-containing proteins. These proteins sense and respond to increasing cellular Pi levels through binding of inositol pyrophosphate (PP-InsP) signaling molecules to their SPX domain^5,6^. XPR1 is the only mammalian protein that contains a SPX domain, and consistent with this, its Pi transport function is a focal point for overall homeostatic maintenance of Pi balance in humans^2,7–9^. A homolog in *Drosophila*, PXo, was also recently shown to play an important role in Pi homeostasis, where it marks multilamellar organelles termed PXo bodies, which are degraded following Pi starvation^10^. Several studies have demonstrated that *in vivo* XPR1 facilitated Pi export requires the PP-InsP, 1,5-bis-diphosphoinositol 2,3,4,6-tetrakisphosphate (1,5-InsP_8_ or more simply InsP_8_), and that other PP-InsPs, such as InsP_6_, 1-InsP_7_, and 5-InsP_7_ have no effect on its activity – clarifying that InsP_8_ is the physiologically-relevant ligand^8,11,12^. The concentration of InsP_8_ in mammalian cells is 100-500 nM with cytosolic levels rising and falling in response to corresponding fluctuations in cytosolic Pi^13–15^. This correlation between cellular Pi and InsP_8_ levels has also been well documented in yeast and plants^16,17^. The mechanism through which the SPX domain of XPR1 discriminates between PP-InsP metabolites is not well understood. Isolated SPX domains have been shown to interact with negatively charged sulfate (SO_4_^2-^) and PP-InsPs with minimal specificity and selectivity^5,8,11^. While the binding affinity of InsP_8_ towards the isolated SPX domain of XPR1 is higher than for other PP-InsPs^8^, given their cellular abundance, these differences in affinity are not sufficient to explain why only InsP_8_ supports Pi export through XPR1 *in vivo*^18^.

Highlighting critical roles in human physiology, aberrant XPR1 transport activity is associated with both brain disease and cancer. Loss-of-function mutations in XPR1 lead to primary familial brain calcification (PFBC), an inherited neurological disorder in which the formation of abnormal calcium phosphate deposits is causative of neuropsychiatric, cognitive, and movement disorders^19–22^. Clinical features include ataxia, Parkinsonism, psychosis, dementia, and cognitive dysfunction with diverse severity, even within the same family.

Furthermore, these deleterious calcium phosphate deposits are also prevalent in ∼20-30% of older adults with no genetic risk factor^23^. Also of importance for human health and disease, highly proliferative cells, such as cancer cells, have increased metabolism and high ATP and Pi needs. Overexpression of the Pi importer, SLC34A2, is a hallmark of ovarian and uterine cancers^24,25^. Preventing Pi export from these cancer cells via genetic or pharmacological XPR1 inhibition leads to Pi toxicity and cell death^26,27^. As such, inhibition of XPR1 is an attractive treatment option for these highly lethal cancers, especially as XPR1 antagonists as a targeted therapy have the potential to minimize side effects.

Despite the importance of Pi homeostasis in human physiology, we have an incomplete understanding of the complex cellular mechanisms that govern Pi entry into and efflux from cells. To obtain a molecular-level understanding of the mechanisms of Pi export through XPR1 and their regulation by elevated cellular InsP_8_ levels, we determined a series of single-particle cryo-electron microscopy (cryo-EM) structures of human XPR1 in distinct conformational states. This includes structures in an apo form and in complex with the substrate, Pi, and the physiologically relevant signaling molecule, InsP_8_. The structures and complementary functional studies reveal the mechanisms of activation by InsP_8_ orchestrated through its regulatory intracellular SPX domain. This work provides a framework for understanding XPR1 function, its transporter cycle, its gating by InsP_8_, and how activity is modulated.

### Functional characterization of XPR1

To allow for the measurement of Pi efflux through XPR1, we developed a cellular radioactive [^32^P] transport assay, and measured a robust signal in hTERT RPE-1 cells (Figure 1A). RPE-1 cells have several benefits. They have a normal karyotype, are non-tumorigenic, and have superior plate adherence characteristics, thereby decreasing nonspecific Pi release. To verify that the Pi efflux that we observed is dependent on endogenous XPR1, we generated XPR1 CRISPR/Cas9 knock-out cells and used a previously characterized inhibitory peptide referred to as XRBD^4^. XPR1 is a cell-surface receptor for xenotropic and polytropic murine leukemia retroviruses (X- and P-MLV), and XRBD is a soluble ligand generated from the envelope-receptor-binding domain of X-MLV that is capable of inhibiting Pi export through XPR1. Export activity was minimal in XPR1 knock-out cells or when the cells were incubated with XRBD (Figure 1A). Next, we stably reintroduced wild-type XPR1 to the knockout cells and measured export of radiolabeled Pi. Pi export function is rescued in these cells (Figure 1B). Though there is evidence that XPR1 can also localize to intracellular organelles^9^, these results confirm that functional XPR1 resides in the plasma membrane of these cells.

**Figure 1.**
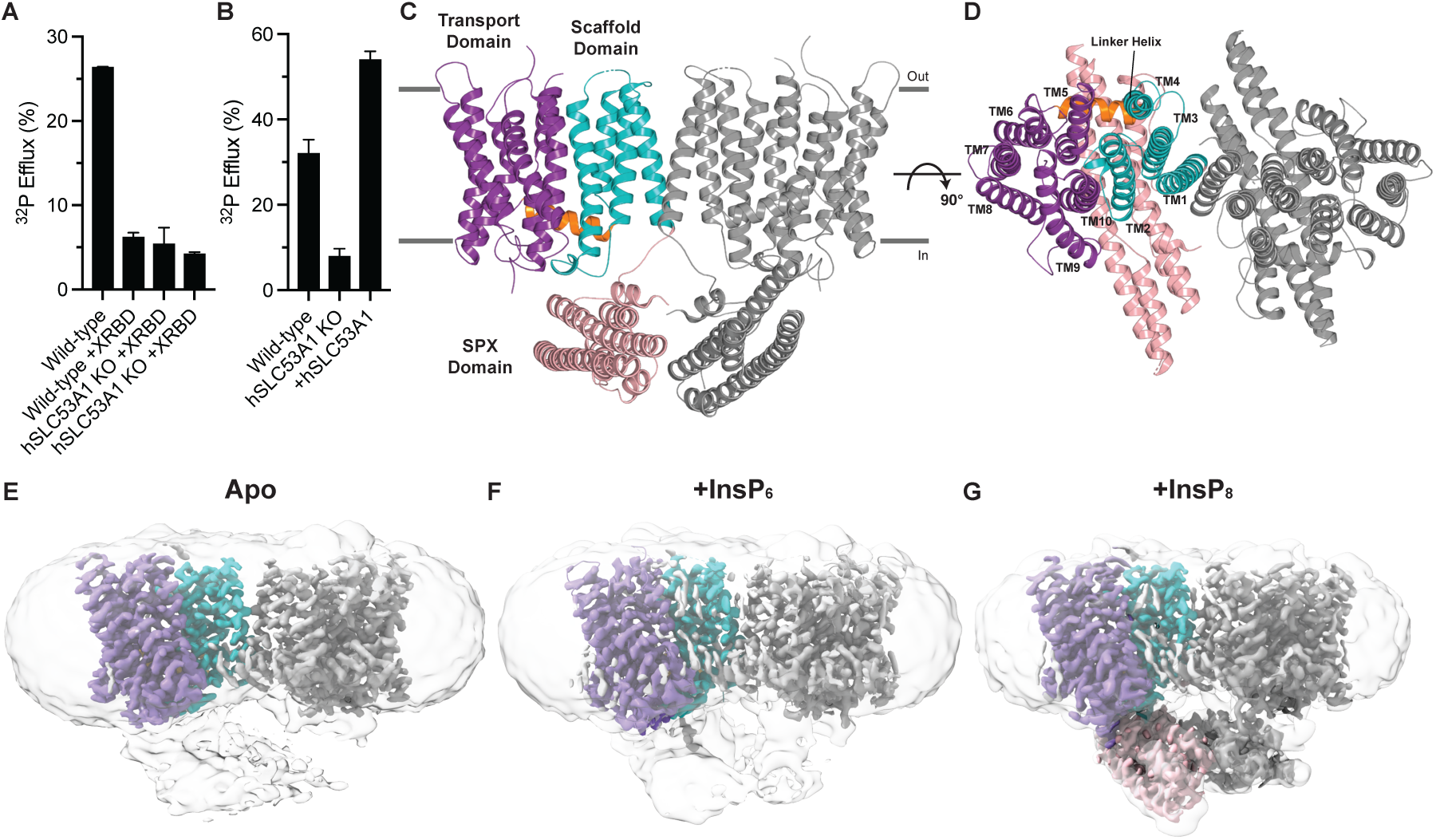
Functional characterization and structures of human XPR1. **(A)** Pi export through endogenous XPR1 in hTERT RPE-1 XPR1-knockout cells, and its inhibition by the viral peptide fragment XRBD. Data are represented as the mean ± SD (n=3). **(B)** Pi export capabilities were regained when wild-type XPR1 was reintroduced into the XPR1-knockout background. Data are represented as the mean ± SD (n=3). (**C-D)** Cartoon representation of the XPR1 structure with InsP_8_ bound and the SPX domain resolved, viewed parallel to the membrane plane (**C**) and perpendicular to the membrane from the extracellular space **(D)**. The SPX domain is colored pink, the scaffold domain is colored teal, the linker domain is colored orange, and the transport domain is colored purple. Horizontal lines indicate approximate boundaries of the plasma membrane. **(E-G)** Cryo-EM density maps of the XPR1 dimer, one monomer colored identically to panels **(C-D)** and the other monomer colored grey, in the absence of PP-InsPs **(E)** and the presence of InsP_6_ **(F)** or InsP_8_ **(G)**. Maps with a low-filter threshold of 6 Å are overlaid.

### Architecture of XPR1

Recombinant full-length human XPR1 protein was purified in the detergent lauryl maltose neopentyl glycol (LMNG) and supplemented with glycol-diosgenin (GDN). The GDN was necessary to overcome preferred orientation of the protein on the cryo-EM grids. We determined single particle cryo-EM structures of XPR1 in distinct conformation states, and also when supplemented with substrate (Pi) and two PP-InsP signaling molecules (InsP_6_, which lacks biological activity, and InsP_8_ that we synthesized^28^, the physiological activator). The average resolutions of our structures range from 2.5 to 3.3 Å. The cryo-EM density maps are of sufficient quality to allow for construction of atomic models of the protein and its corresponding substrates and ligands with good stereochemistry and correlation with the density (Table S1 and Figure S6).

The overall architecture of XPR1 will first be described for our samples incubated with InsP_8_ prior to structure determination, as these maps are the most complete. The structure of XPR1 is comprised of an N-terminal SPX domain and ten transmembrane ɑ-helices (TM1 to TM10) – the majority of the C-terminal tail (residues 633-696) is disordered in all our cryo-EM maps (Figure 1C,D). XPR1 assumes a dimeric conformation. The membrane-embedded domain of the protein can be divided into two regions: the scaffold domain formed by TM1-4, which permits dimerization, and the transport domain formed by TM5-10, where the substrate translocation pathway lies. The two domains are connected by a helix that runs parallel to the inner leaflet of the membrane designated the linker helix. We detected particularly striking densities for a cholesterol molecule and a phosphatidyl lipid at the interface between these two domains (Figure S8A). The cholesterol molecule that we observed is lodged between aromatic residues (Tyr558 and Phe609) of TM9 and TM10 (Figure S8B). The phosphatidyl lipid occupies an internal cavity formed by TM2, TM3, TM4, the linker helix, and TM5 (Figure S8C).

### Visualizing XPR1 in distinct conformational states

We obtained three distinct conformational states of XPR1 that are correlated with the presence and identity of PP-InsPs. In our InsP_8_-bound structures, the intracellular SPX domain, which is thought to bind InsP_8_ and is essential for XPR1 function, is well ordered (Figure 1G)^9^. These maps stand in stark contrast to those generated from the apo and InsP_6_-supplemented protein where this intracellular sensing domain cannot be resolved (Figure 1E,F). By low-pass filtering these maps, we can conclude that the SPX domain is present, but that in the absence of the physiological activator, InsP_8_, this region of XPR1 is highly dynamic. The movements are likely imparted by the flexible linker (residues 223-229) that connects the SPX and transmembrane domains.

When we supplemented high Pi (50 mM – physiological concentrations are ∼1-2 mM) in addition to InsP_6_, we resolved a third state, following extensive particle classification, in which the SPX domain is resolved (Figure S9). The map quality for the SPX domain is poor, indicating that there is large degree of heterogeneity in this region. However, bound InsP_6_ is visible. This conformation is characterized by an asymmetric SPX dimer that is positioned beneath the transmembrane domain of one of the monomers. We are doubtful that this conformation exists in cells. Our results suggest that non-physiological concentrations of Pi can influence XPR1 protein behavior, perhaps acting in concert with PP-InsPs.

Comparison of our XPR1 maps suggests that InsP_8_ is altering the dynamics of the SPX domain. A key finding is that InsP_6_, though similar in chemical structure, is not capable of recapitulating the stabilizing effect of InsP_8_. Structures of a nonhomologous yeast Pi importer that contains a SPX domain have been determined using cryo-EM, and revealed that its SPX domain is highly dynamic^29^. Furthermore, NMR indicates that movents of the SPX2 domain of the yeast heterooligomeric vacuolar transport chaperone (VTC) complex are altered by PP-InsP binding^30^. This further suggests that the dynamic nature of SPX domains may be a conserved feature and may be key to their mechanisms of action, even in proteins with diverse functions.

### Dimerization is essential for XPR1 function

Our structures reveal that XPR1 is a homodimer, with dimerization being facilitated by a GxxxG ɑ-helical packing motif located in TM1 (Figure S10A)^31^. To determine the importance of the residues in the GxxxG motif, we mutated the two glycine residues to leucine and employed our previously described cellular radioactive transport assay. Our results demonstrate that this dimerization is essential for the proper transport function of XPR1 (Figure S8B).

Whereas this dimer population is the most prevalent, we did observe an alternate dimer assembly, in which one of the monomers (TM2-10) appear to be rotated via a large rigid-body movement with respect to the other (Figure S10C). Overlay of these structures reveals that the rotation is caused by a large rearrangement of TM1 (Figure S10D). Cholesterol is known to bind, stabilize, and modulate the activity of several transporters^32^. The cholesterol molecule that we observed copurified with XPR1 is absent when TM1 assumes this alternate position, suggesting that cholesterol may play a role in stabilizing TM1 and permitting dimerization and XPR1 function (Figure S10E). In its place, we observed a density for a phospholipid, whose presence implies that this hydrophobic pocket must be filled. Though the implications of our findings require further investigation, they suggest that dimerization may serve as a regulatory mechanism of XPR1 function, one in which cholesterol binding may play a modulatory role.

### Substrate transport pathway

In our XPR1 maps, we observed three discrete densities in the transport domain that are consistent with Pi substrate (Figure 2A). To facilitate the confident assignment of these densities as Pi ions and the identification of additional binding sites, we determined structures in the presence of a physiological concentration (5 mM) of Pi. The same three densities were present, and no others were identified. The strength of these densities varies between maps; however, it is apparent that Pi at least partially occupies these sites even when it is omitted during protein purification. This suggests that the nominal Pi in our expression and purification buffers is sufficient to populate the Pi-binding sites and implies high-affinity binding.

**Figure 2.**
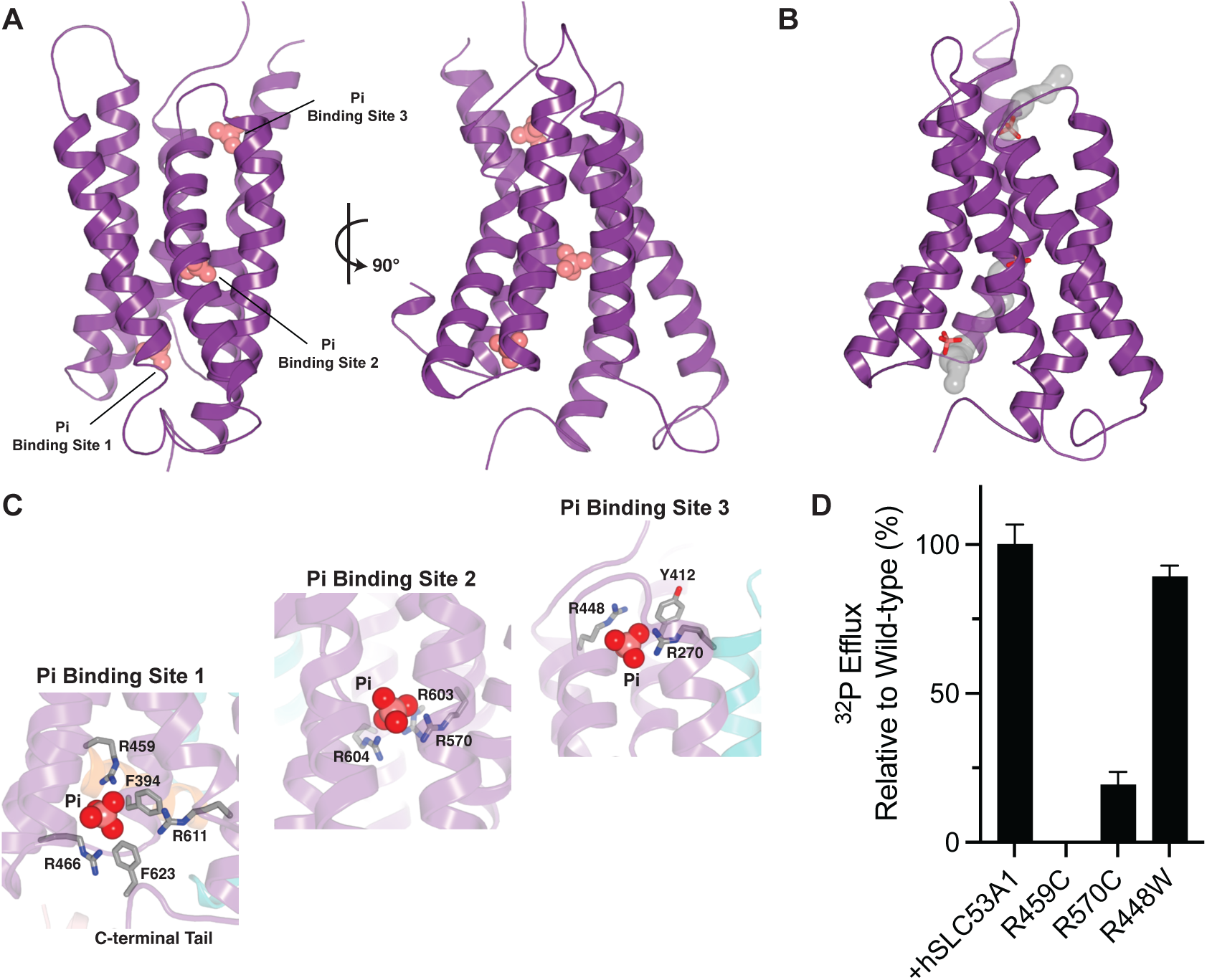
The substrate translocation pathway. **(A)** The position of the three Pi ions (pink spheres) helps define the substrate translocation pathway. (**B)** Tunnels leading from the cytosol to Pi sites 1 and 2 and from Pi site 3 to the extracellular space. There is no direct path between Pi sites 2 and 3 in this conformational state. (**C)** Interactions with the Pi substrate in the three bindings sites. Nitrogen, blue; oxygen, red. (**D)** Export of radioactive ^32^P by XPR1-knockout RPE-1 cells stably expressing wild-type or mutant XPR1-mCer. These mutants are found in patients with PFBC. Data are represented as the normalized mean ± SD (n=3).

The position of the Pi-binding sites defines the substrate transport pathway through XPR1. Substrate to be exported from cells first encounters site 1 – the substrate-binding site closest to the inside of the cell (Figure 2B). Site 1 is connected to site 2 via a short tunnel and lies approximately at the mid-way point of the membrane-embedded substrate translocation pathway. The final site, site 3, lies adjacent to the extracellular space. The lengthy tunnel connecting Pi sites 2 and 3 is blocked in this conformational state by Trp573, suggesting that rearrangement would be required for substrate in sites 1 and 2 to reach site 3 and the outside of the cell. Direct binding of negatively charged Pi is commonly promoted through positive electrostatic potential. Indeed, all three Pi ions are coordinated by two or more positively charged residues (site 1: R459, R466, R611; site 2: R570, R603, R604, site 3: R270, R448) (Figure 2C). Several of these Arg residues are positioned via cation-π interactions with Tyr and Phe residues.

Mutation of amino acid residues in these binding sites are known to cause the neurological disorder, PFBC, due to a lack of function of XPR1 (Figure 5)^33–35^. To ascertain the functional relevance of the interactions observed, we stably introduced three disease-causing mutations (R459C at site 1, R570C at site 2, and R448W at site 3) into our XPR1 CRISPR knock-out cell line, and measured Pi export activity. While the R448W mutation was tolerated, R459C and R570C have reduced transport function – R459C is nonfunctional and R570C has limited activity (Figure 2D). All three mutants had fluorescence-detection size-exclusion chromatography (FSEC) profiles that were nearly identical to the wild-type transporter (Figure S11A), indicating that they did not markedly affect protein expression, folding, or stability despite the dramatic residue changes. This indicates that Pi binding to the sites adjacent to the intracellular environment, site 1 and to a lesser extent site 2, is essential for substrate translocation. It also suggests that substrate bind to the site proximal to the extracellular space, site 3, plays a minimal role in XPR1 function. Therefore, the ability of Pi to enter the substrate translocation pathway is of upmost importance for productive transport.

### Structural mechanisms of InsP_8_-dependent regulation

To determine the structural changes and molecular mechanisms governing InsP_8_-mediated activation of XPR1, we further analyzed our structures resulting from protein prepared in the presence of InsP_8_ (Figure 3A). As was previously discussed, in these maps the SPX domains have been rigidified, and the resolution is sufficient in this region for reliable modeling of both protein and ligand. Our structures reveal that four InsP_8_ molecules bind the XPR1 dimer at two unique sites (Figure 3B). In these sites, the highly negatively charged InsP_8_ is embedded within surface pockets of positive electrostatic potential.

**Figure 3.**
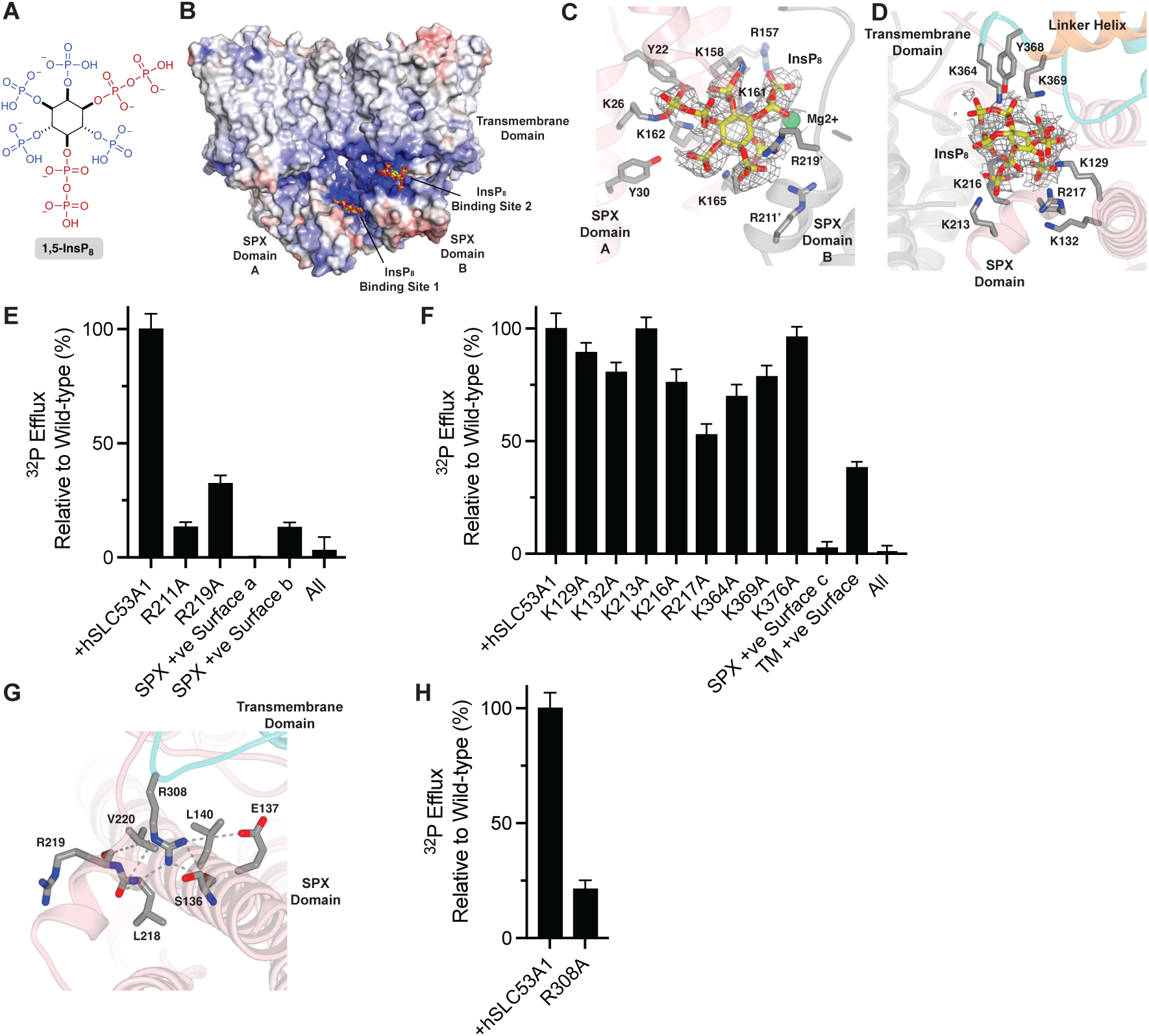
Structural basis of InsP_8_ recognition by XPR1. **(A)** Chemical structure of 1,5-InsP_8_. The characteristic pyrophosphate groups are highlighted in red. **(B)** XPR1 contains two positively charged pockets in which InsP_8_ binds forming a bridge between neighboring domains and locking the transporter in a defined conformation with rigid SPX domains. The molecular surface is colored according to electrostatic potential: light gray regions are neutral; red, −8 kTe^-1^; blue, +8 kTe^-1^. **(C-D)** Close-up view of InsP_8_-binding site 1 **(D)** and InsP_8_-binding site 2 **(D)** with InsP_8_ densities (gray mesh, 5σ contour **(C)** and 3σ contour **(D)**). Nitrogen, blue; oxygen, red. **(E-F)** Structure-function analysis demonstrating the importance of the positively charged surface patches that bind InsP_8_ for transport function (binding site 1 **(E)** and binding site 2 **(F)**). Data are represented as the normalized mean ± SD (n=3). **(G)** Interactions between the Arg extension contributed by the transmembrane domain and the intracellular capture pocket. Dash lines indicate electrostatic interactions. Nitrogen, blue; oxygen, red. (**H)** Structure-function analysis elucidating the import of Arg308, and its key role in stabilizing the SPX domain and the inward-open state of the transporter. Data are represented as the normalized mean ± SD (n=3).

The first InsP_8_ binding site is comprised of two binding interfaces, one contributed by monomer A and one contributed by monomer B (which we refer to as SPX +ve surface a and SPX +ve surface b, respectively) that form a pocket between the two neighboring SPX domains (Figure 3C). SPX +ve surface a is comprised of Tyr22, Lys26, Tyr30, Arg157, Lys158, Lys161, Lys162, and Lys165 (Figure 3C), and it is known to be functionally important for binding of negatively charged moieties to SPX domains, such as SO_4_^2-^ and InsP_8_, as well as InsP_8_-dependent Pi homeostasis^5,9,36^. SPX +ve surface b is formed by Arg211 and Arg219. We also observed a coordinating metal ion that we have modeled as magnesium (Mg^2+^). Binding of InsP_8_ to this site stabilizes a SPX dimer by bridging this domain between monomers. This dimer configuration was not observed in the previous crystal structures of the SPX domain alone with InsP_8_ as the construct was truncated at residue 207 and therefore was missing SPX +ve surface b^9^. The importance of the 1- and 5-diphosphate groups of InsP_8_ for binding to site 1 is apparent (Figure 3C): the diphosphates make critical polar contacts with XPR1, including for one pyrophosphate – Arg157 and Lys158 and for the other pyrophosphate – Lys26, Lys162, and Tyr22, as well as the backbone nitrogens of Lys2, Phe3, and Ala4.

The second InsP_8_ binding site is positioned both by an additional set of conserved residues within the intracellular SPX domain (which we designate as SPX +ve surface c) and a surface patch of positively charged residues contributed by the intramembrane scaffold and linker domains (referred to as the TM +ve surface) (Figure 3D). Specifically, InsP_8_ at this site engages in electrostatic interactions with Lys129, Lys132, Lys213, Lys216, and Arg217 of SPX +ve surface c and Lys364 and Lys369 of the TM +ve surface. Residue Tyr368 likely positions Lys364 and Lys369 via cation-π interactions. The density for this second InsP_8_ molecule is weak in comparison to both the surrounding protein and the InsP_8_ molecule bound to site 1. We attribute this to the pocket being cavernous and able to accommodate numerous poses of InsP_8_. Of note, the phosphatidyl lipid that we found fills an internal cavity has its headgroup positioned nearby to the InsP_8_-binding site (Figure S8C). Ultimately, binding of InsP_8_ to this site helps to bridge the SPX and transmembrane domains further reducing the dynamics of the SPX domain.

To determine whether the two binding sites that we have identified are responsible for InsP_8_-dependent activation, we analyzed Pi export for XPR1 transporters bearing single alanine substitutions at several of the identified InsP_8_-binding residues. The importance of the InsP_8_-binding site 1 interface on SPX +ve surface a has been previously appreciated^5,36^. To evaluate whether SPX +ve surface b is also critical, we mutated these arginine residues. Indeed, mutants R211A and R219A show diminished transport function, validating their critical role in the binding of InsP_8_ to the SPX domain of XPR1 and the importance, not just of a single positively charged surface, but of an entire positively charged pocket (Figure 3E). Given the import of InsP_8_ binding to site 1 for proper transport function, we speculate that dimerization of the XPR1 transmembrane domain is essential because it brings the two interfaces that comprise the binding pocket into close proximity. For InsP_8_-binding site 2, single point mutations within the positively charged clusters on either surface had little effect on transport activity (Figure 3F). This suggested to us that the major determinant of ligand recognition by this second site is the overall positive charge within these surfaces and the pockets they create.

To test this idea, we analyzed mutants lacking all positive charges within these pockets. For InsP_8_-binding site 1, this includes SPX +ve surface a (K26A, R157A, K158A, K161A, K162A, K165A) and SPX +ve surface b (R211A, R219A). For InsP_8_-binding site 2, this includes SPX +ve surface c (K129A, K132A, K213A, K216A, R217A) and TM +ve surface (K364A, K369A, K376A). We found that these XPR1 mutants can no longer or very minimally export Pi, nor can mutants in which all positive charges in the binding sites are converted to alanine (Figure 3E,F). Together, these data indicate that binding of InsP_8_ to both pockets is necessary for XPR1 transport function and that all four of the electropositive surfaces that we have identified are important for InsP_8_ binding. Furthermore, our results demonstrate that InsP_8_ binding is not wholly intrinsic to the SPX domain of XPR1 but requires interactions with the neighboring transmembrane domain.

Further emphasizing the importance of the transmembrane domain of XPR1 in the InsP_8_ activation mechanism, our maps reveal that Arg308 extends from an intracellular loop of the transmembrane domain down into the SPX domain where it is positioned in a pocket to enhance stability and help lock the SPX domain in this distinct state (Figure 3G). This interaction is strengthened by electrostatic interactions involving Glu137 and the backbone carbonyl groups of Ser136, Leu218, and Arg219. The hydrophobic residues Leu140 and Val220 also help define the pocket shape. Arg308 is critical for XPR1 transport activity (Figure 3H). Three known PFBC associated mutations – S136N, L140P, and L218S are localized to the Arg capture pocket in the SPX domain and have a documented reduction in function (Figure 5)^22^. Notably, in our structures where the SPX domain is disordered (apo or InsP_6_-supplemented), the Arg308 side chain is also not visible.

Communication between the two InsP_8_ sites and the Arg extension may be facilitated through the linker (residues 212-230) that connects the SPX domain to TM1. Specifically, Arg211 and Arg219 help form InsP_8_ site 1, whereas Lys213, Lys216, and Arg217 help form InsP_8_ site 2, and Leu218, Arg219, and Val220 help form the Arg extension and capture pocket.

The identification of how InsP_8_ binds provides additional support that it is the dynamics of the SPX domain that are crucial for the InsP_8_-dependent regulatory mechanisms of XPR1. Furthermore, they indicate how InsP_8_ locks the SPX domain into a single state. At physiological pH, InsP_8_ is highly negatively charged (−10 or −11)^37^, which allows it to interact with more than one positively charged protein site simultaneously. This allows for the bridging of surfaces that would otherwise repel each other. Our structures indicate that InsP_8_ binding at interfaces between the SPX monomers and SPX and transmembrane domains restricts movement of the intracellular SPX domain and stabilizes a defined transporter state. Together, these observations illustrate how InsP_8_ co-factor binding significantly alters protein behavior.

### InsP_8_-induced movements initiate the transport cycle

Further classification of particles originating from the InsP_8_ supplemented samples yielded two different states along the transport cycle. In both states the SPX domains are engaged and InsP_8_ is bound. The superposition of these states revealed that rearrangement of the C-terminal tail originating at His616 is a hallmark of the transition from an inward-open to an occluded conformation (Figure 4A). Highlighting the importance of this region, the C-terminal tail residues of XPR1 are highly conserved (Figure S7). We observed dimers in which one protomer is in an inward-open state and the other in an occluded state and in which both protomers adopt the occluded state. As monomers have independent substrate translocation pathways and both XPR1 states can be visualized within the dimer, the protomers appear to have low cooperativity.

**Figure 4.**
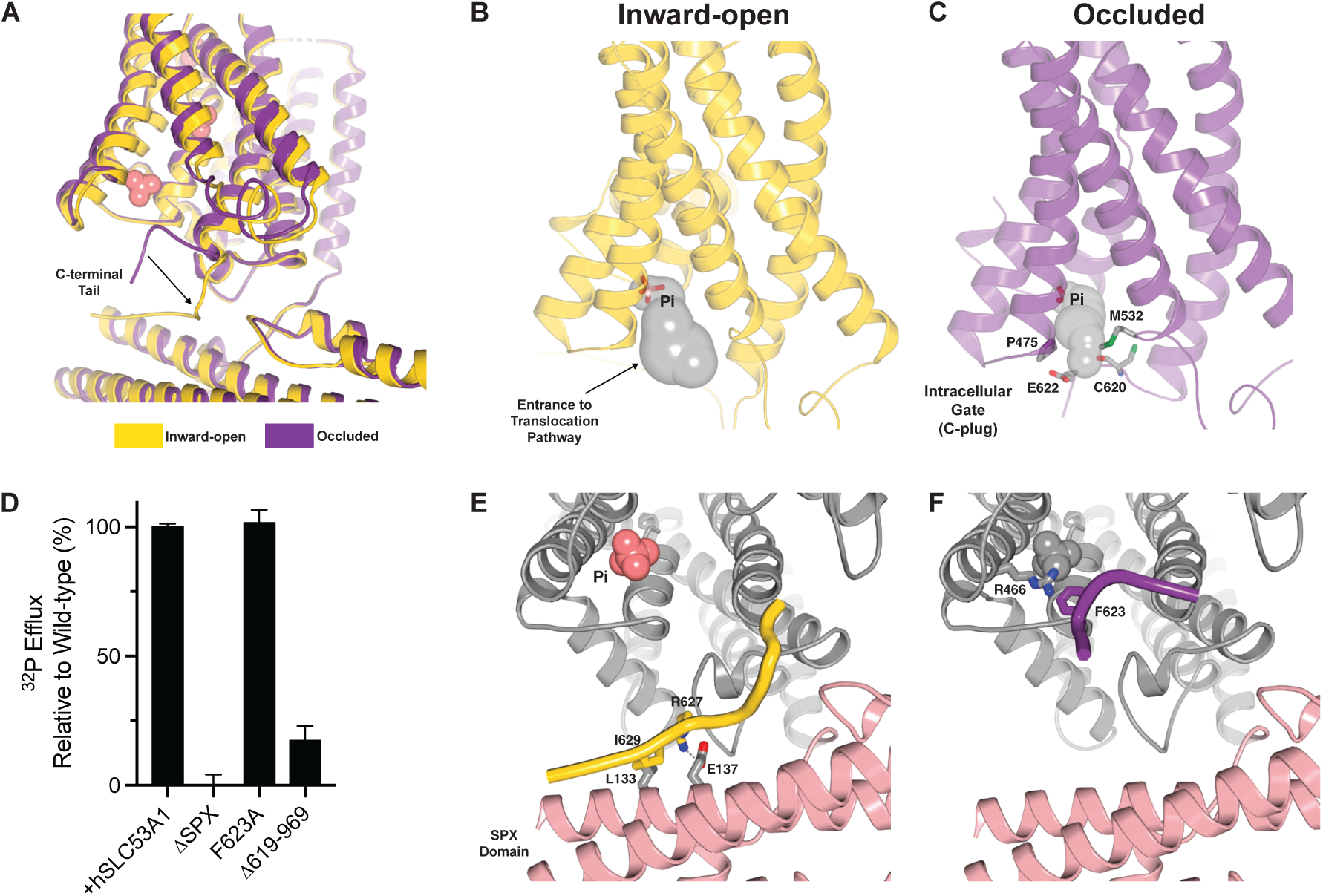
InsP_8_ binding opens the intracellular gate. **(A)** Superimposition of the inward-open (yellow) and occluded (purple) XPR1 structures. **(B-C)** The C-terminal fragment (C-plug) of XPR1 blocks an otherwise inward-open conformation. The intracellular gate is formed by Pro475, Met532, Cys620, and Glu622. Contour mesh of the tunnels (gray) calculated by MOLE. **(D)** Structure-function analysis demonstrating the importance of the intracellular domains on XPR1 transport activity. Data are represented as the normalized mean ± SD (n=3). **(E-F)** Structural changes associated with XPR1 gating. Transition from the inward-open **(E)** to occluded **(F)** state is accompanied by an insertion of a portion of the C-terminal tail of XPR1 into the entrance to the Pi transport pathway. This forms the intracellular gate.

In the occluded state, the C-terminal tail residues Cys620 (via its backbone carbonyl) and Glu622 block Pi entry to the translocation pathway by introducing negative charge that repels the Pi ions and restricts its size (Figure 4C and S12B). A neighboring glycine residue, Gly621, also plays a critical role as it allows these residues to assume an optimal conformation. We define this region as the intercellular gate or C-plug. This C-terminal tail position is stabilized through a cation-π interaction of Phe623, which projects into Pi-binding site 1, with Arg466 – one of the Pi coordinating residues (Figure 4F). This conformation predicts that substrate binding to site 1 is necessary to drive rearrangement from the inward-open to occluded state.

In contrast to the occluded XPR1 structure, our inward-open structure shows an unplugging of the entrance of the Pi translocation pathway through a sequestering of the C-terminal tail by direct interactions with the SPX domain. This increases the radius of the entrance to the translocation pathway from 1.6 Å to 4 Å, which is large enough to allow passage of a Pi ion (radius ∼3 Å) (Figure 4B). Furthermore, the entrance to the Pi permeation pathway is lined by residues Lys388 and Arg472 and thus is highly electropositive (Figure S12A). A critical salt bridge is formed between Arg627 of the C-terminal tail and Glu137 of the SPX domain and a series of hydrophobic interactions are also contribute to this interaction (Figure 4E). The importance of this C-terminal tail region is further emphasized by the nearby location of several PFBC associated mutations, including N619D, R624H, and I629S (Figure 5)^33,38^. Worth noting, this key structural feature neighbors the interactions of the Arg308-containing intracellular loop that helps bridge the SPX and transmembrane domains.

**Figure 5.**
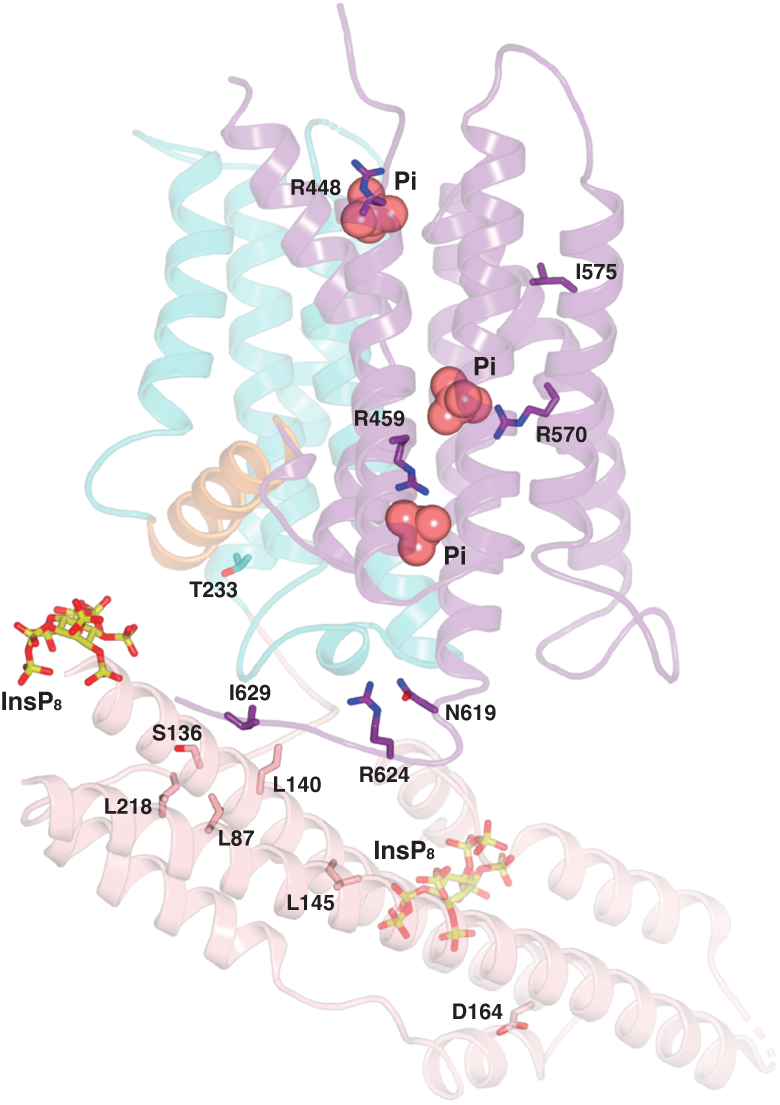
Locations of PFBC disease mutations mapped on the inward-open structure of XPR1. Residues of which missense mutations are known to cause neurological disease phenotypes are represented as sticks. Nitrogen, blue; oxygen, red. The residues are clustered near protein regions critical for function, including 1) the substrate translocation pathway, 2) the Arg extension and capture pocket, and 3) the C-terminal tail.

Our structures predict that in the absence of a SPX domain (and InsP_8_), XPR1 will be unable to enter the inward-open state and will therefore be nonfunctional. Indeed, all our structures from samples without InsP_8_, whether the SPX domain is resolved or not, are in an occluded state. Reports on whether the SPX domain is essential for function have been conflicting^4,9,36,39,40^. In contrast, the importance of the C-terminal tail is clear since deletion of this domain (Δ612-696) renders XPR1 non-functional – although this may be attributed in part to protein misfolding and/or mislocalization^33^. To further assess the role of these intracellular domains on transport, we generated mutant transporters lacking the SPX domain (Δ1-225), lacking the C-terminal tail (Δ619-696), or bearing a single substitution of Phe623, the C-terminal tail residue that participates in substrate binding in the occluded state, to alanine. We expressed the mutants in XPR1-knockout cells and measured export of radiolabeled Pi. Transporter lacking the SPX domain was completely non-functional, indicating that this protein region is, indeed, essential for Pi transport (Figure 4D). Consistent with previous studies, removing the C-terminal tail of XPR1 had a detrimental effect on protein behavior. The lack of a FSEC peak for this mutant suggests that it is either not expressed or that the expressed protein is not stable following detergent extraction (Figure S11E). The function of the F623A substitution was comparable to wild-type transporter, though the FSEC peak is broader, implying that there may be increased conformational heterogeneity.

The mechanism of XPR1 activation is reminiscent of several other transporter families that conduct anions and for which their C-terminal tails are crucial for the transport cycle. For example, in the occluded state of the glutamate/gamma-aminobutyrate (GABA) antiporter GadC, the C-terminal tail is folded within the open binding cavity, thus blocking the substrate-binding site^41^. In SLC26A9, another transporter that mediates anion export activity, occupation of the intracellular pocket by its C-terminal tail leads to a change in electrostatics at the entrance of the translocation pathway and reduces accessibility from the cytosolic side, thereby altering substrate access^42^.

Given the detailed mechanisms of InsP_8_ activation and our structures of InsP_6_-supplemented protein, it is now apparent why InsP_6_ is unable to activate XPR1 transport function *in vivo*. At physiological concentrations of Pi, InsP_6_ is not capable of stabilizing the SPX domain, a necessity for conversion to the inward-open state and Pi export. It is plausible that InsP_6_ binds, though it is not possible to say with certainty due to the dynamics of the SPX domain and corresponding poor map quality. Regardless, it is apparent that InsP_6_, which has a lower negative charge than InsP_8_, cannot facilitate robust domain bridging. At extremely high concentrations of Pi (∼20X cellular levels), we did obtain InsP_6_-bound structures where the SPX domain is resolved, albeit with poor resolution (Figure S9). The SPX dimer has an alternate placement relative to the transmembrane domains and adopts this asymmetric conformation because InsP_6_ binding is not analogous to InsP_8_ binding. We observed only two InsP_6_ molecules bound to the XPR1 dimer. One InsP_6_ binds only to SPX +ve surface a of monomer A. In the absence of a second interface, domain bridging is not possible. The second InsP_6_ binds to the SPX +ve surface a of monomer B and the TM +ve surface of monomer A, explaining why the position of the SPX domain relative to the transmembrane domain is different than what we observed for InsP_8_-bound XPR1. Though the dynamics of the SPX domain have been altered, the pose of this domain is not capable of releasing the C-plug and cycling the transporter to the inward-open state.

Our structures support a mechanism for the dependence of XPR1 function on InsP_8_ as depicted in Figure 6. The intracellular regulatory gate (or C-plug) is formed by the C-terminal tail of XPR1. Upon InsP_8_ binding, the SPX domain becomes rigidified in a conformation that can sequester the C-terminal tail of XPR1, causing it to undergo a dramatic rearrangement away from the entrance of the translocation pathway. This leads to opening of the intracellular gate. When open, Pi ions are free to enter the substrate translocation pathway and bind at sites 1 and 2. We propose that the plugging of the entrance of the translocation pathway by the C-terminal tail is a movement that is favored by the participation of Phe623 to substrate binding to site 1 and may help initiate the transition to the occluded state. Even with InsP_8_-bound and the SPX domain engaged, the majority of our particles (∼75%) reside in the occluded state. This may be because in our strutures, despite not introducing exogenous Pi, we find substrate binding site 1 is occupied. Lastly, our structures predict that a conformational change to the outward-open state, substrate release, and completion of the transport cycle, requires relieving two points of constriction along the ion conduction pathway. This includes the constrictions caused by Trp583, which is located mid-way through the substrate tunnel, and an extracellular gate clearly present in our inward-open and occluded conformations. While both the TM5-6 and TM9-10 loops occlude the release of Pi from substrate binding site 3, we hypothesize that the extracellular gate is formed by the TM9-10 loop, as the upper segments of TM5 and TM6 are stabilized by a disulfide bond between cysteines 415 and 440.

**Figure 6.**
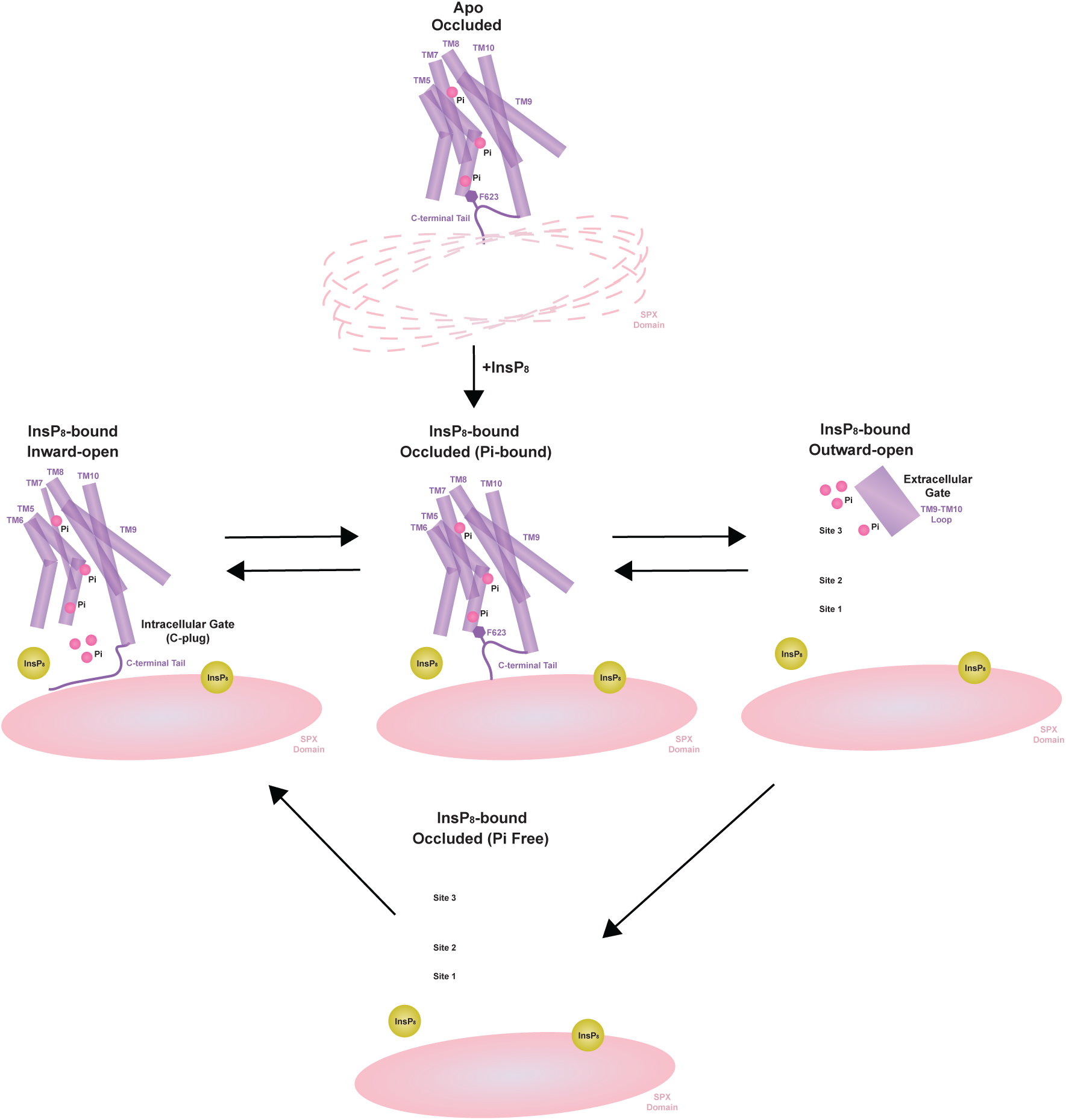
Transport cycle of SLC53A. In the presence of InsP_8_, the SPX domain becomes rigidified and the C-terminal tail of XPR1 is sequestered by this domain causing a transition to an inward-open state. Repositioning of the C-terminal tail and insertion of Phe623 into substrate-binding site 1, represents the transition to an occluded state. Our structures predict an outward-open state in which a proposed extracellular gate, formed by the TM9-10 loop, must be repositioned to allow for Pi release.

## Discussion

### Structural basis of alternating access of XPR1

**–** Transporters shuttle molecules across cell membranes by alternating among distinct conformational states. Three major states, inward-open, occluded, and outward-open, allow the substrate to access the transport pathway from both sides of the membrane. These global structural transitions are a necessary means of information transfer between the two sides of a transporter. Our structures suggest that XPR1 is a Pi uniporter that facilitates passive diffusion down the concentration gradient. This is in contrast to Pi importers which are Na^+^-dependent^3^. Our structures also demonstrate that XPR1 has multiple substrate-binding sites, and it will be interesting to determine the stoichiometry of the transport cycle. In the absence of InsP_8_ and at physiological Pi concentrations, the resting state of the intracellular gate of the transporter is closed. Binding of InsP_8_ triggers a conformation transition to the inward-open state. Upon binding of Pi to sites 1, a further conformational change occurs that captures Pi within the transporter and defines the occluded state. Subsequently, another rearrangement must occur to produce the outward-open state so that substrate molecules can be released. We have identified three gates: 1) an intracellular gate, formed by the C-terminal tail, 2) a gate within the substrate translocation pathway between Pi binding sites 2 and 3 in the inward-open and occluded states, and 3) an extracellular gate, likely formed by the TM9-10 loop.

### Diversity of mechanisms involving SPX domains

– XPR1 is the only mammalian protein known to sense Pi concentrations through the binding of PP-InsPs to adjust its activity to meet the needs of the cell. Intriguingly, to maintain cell Pi homeostasis, it has been suggested that Pi uptake and Pi efflux must be coordinated^8,9,36^. Though InsP_8_ has emerged as a central cellular messenger, our understanding of the molecular mechanisms that govern protein regulation by this signaling molecule is limited^43^. A major advance was the revelation that InsP_8_ binds to and signals through SPX domains^5^. Our results demonstrate that when XPR1 binds InsP_8_, this signaling molecule aids in bridging domains, thereby stabilizing the SPX domains and an active state of the transporter. The ability of PP-InsPs to act as a bridging moiety between positively charged protein surfaces that would otherwise repel each other is well documented – though this is the first example to our knowledge of InsP_8_ acting as a molecular glue^44,45^. The binding pockets for the two InsP_8_ molecules are formed by four positively-charged surfaces – only one of which had been previously identified through studies conducted with the isolated SPX domain^5^. We have shown that InsP_8_ binds not only the SPX domain, but also to the transmembrane-embedded domain of XPR1. This reveals that other protein domains, in addition to the SPX, are important for determining mechanisms of InsP_8_ action. Prior to this work, our molecular-level understanding of how PP-InsP binding stimulates protein function centered on the idea that binding relieves an inhibitory constraint imposed by the SPX domain. For example, in the yeast VTC, biophysical experiments support the notion that InsP_8_ disrupts an interaction between two SPX domains contributed from different proteins within the complex that block inorganic polyphosphate synthesis by the enzyme^30^. Our structures now suggest that modulation of SPX domain-containing proteins by PP-InsPs is mechanistically diverse – with XPR1 being the first example of a protein whose function is regulated by the ability of InsP_8_ to bridge domains and stabilize unique protein states.

### Disease and pharmacological insights

– Over the past decade, six causative genes for PFBC have been identified, two of which are inorganic phosphate (Pi) transporters (SLC20A2 and XPR1), establishing a direct link between the dysregulation of Pi homeostasis and disease^19^. Why these abnormal calcifications form is still yet to be discovered. Furthermore, no disease-modifying therapies are available for brain calcification, nor have definitive pharmacological targets been identified. Our structural elucidation of the Pi exporter XPR1 provides a detailed, mechanistic understanding of how PFBC missense mutations cause impaired Pi efflux and homeostasis and neurological disease. Both agonists and antagonists of XPR1 have potential therapeutic applications, with antagonists being explored as targeted ovarian and uterine cancer drug treatments^26,27^. To address biological questions, and eventually illuminate novel therapies for the treatment of brain calcifications, cancer, and other disorders affected by Pi imbalance, such as kidney disease^46^, access to specific chemical probes that modulate XPR1 transporter function is essential. However, aside from the physiological activator InsP_8_ and the inhibitory viral peptide XRBD, no such compounds currently exist. Our structures suggest that Pi mimics that could block the ion conduction pathway or prevent structural rearrangements during the transport cycle are worth exploring. Our findings also indicate that modulators capable of disrupting the SPX domain stabilization that results when InsP_8_ binds might be effective antagonist drugs. Our findings provide a mechanistic rationale for understanding how Pi transverses the membrane through XPR1 and InsP_8_ initiates XPR1 transporter gating. This may facilitate the design of drugs that selectively modulate aberrant transporter activity under a variety of pathophysiological conditions.

## Data Availability

Cryo-EM density maps have been deposited to the Electron Microscopy Data Bank (EMDB) under the accession numbers EMD-xxxx (apo XPR1), EMD-xxxx (apo XPR1; Alternate dimer); EMD-xxxx (InsP_6_-supplemented XPR1), EMD-xxxx (high Pi- and InsP_6_-supplemented XPR1); EMD-xxxx (InsP_8_-supplemented XPR1; Inward-open/Occluded state), EMD-xxxx (InsP_8_-supplemented XPR1; Occluded state), EMD-xxxx (Pi- and InsP_8_-supplemented XPR1; Inward-open/Occluded state), EMD-xxxx (Pi- and InsP_8_-supplemented XPR1; Occluded state). Atomic coordinates have been deposited in the Protein Data Bank (PDB) under IDs xxxx (apo XPR1), xxxx (apo XPR1; Alternate dimer); xxxx (InsP6-supplemented XPR1), xxxx (InsP_8_-supplemented XPR1; Inward-open/Occluded state), xxxx (InsP_8_-supplemented XPR1; Occluded state), xxxx (Pi- and InsP_8_-supplemented XPR1; Inward-open/Occluded state), xxxx (Pi- and InsP_8_-supplemented XPR1; Occluded state). All DNA constructs and cell lines described in this study are available upon request.

## Acknowledgements

We thank M. Alkareh, L. Cohen-Abeles and J. Luo for their contributions in establishing the radioactive transport assay; R. Hite, D. Julius, C.D. Lima, S. Shuman, and members of the Hite and Diver laboratories for discussions; and M. J. de la Cruz of the Structural Biology core Facility at the Memorial Sloan Kettering Cancer Center for help with data acquisition. M.M.D. is supported by the National Institutes of Health (NIH) National Cancer Institutes (NCI) Cancer Center Support Grant P30-CA008748 and is a Josie Robertson Investigator. H.J.J. is supported by the Deutsche Forschungsgemeinschaft (DFG) under Germany’s excellence strategy (CIBSS, EXC-2189, Project ID 390939984). M.F.Y. receives funds from the NIH/NIGMS through the Weill Cornell Initiative for Maximizing Student Development (IMSD).

## Author Contributions

Q.Z., M.F.Y., and M.M.D. designed and executed experiments, including protein expression and purification, cryo-EM data acquisition and image processing, atomic model building and refinement of XPR1 structures, as well as radioactive transport assays. N.J. and H.J.J. provided the InsP_8_. All authors contributed to preparation of the manuscript.

## Declaration of Interests

The authors declare no competing interests.

## Supplemental Information

Document S1. Figures S1-S12 and Table S1.

## Method Details

### Generation of CRISPR knockout constructs and cell lines

We generated a IL25 knockout hTERT RPE-1 cell line (ATCC, CRL-4000), as a positive control, and a XPR1 knockout RPE-1 cell line. sgRNAs (listed in Table S1) were synthesized by IDT and cloned into pLentiCRISPRv2-mCherry (Addgene, 99154). The plasmid was simultaneously linearized by BsmBI (Fisher Scientific, FERFD0454) and dephosphorylated by Alkalaine Phosphatase (Fisher Scientific, EFREF0651). The sgRNAs were phosphoryated using T4 Polynucleotide Kinase (NEB, M0202S) and annealed in preparation for ligation into the vector (NEB, M2200). The constructs were validated by Sanger sequencing. The sgRNA expressing vectors along with lentiviral packaging plasmid psPAX2 and envelop plasmid pMD2.G-VSVG were transfected into LentiX cells using Lipofectamine 3000 (Thermo Fisher Scientific, L3000015). Virus-containing supernatant was collected 48 h after transfection and passed through a 0.45-μm filter. RPE-1 cells in 6-well tissue culture plates were infected with virus and 4 μg/ml Polybrene (Santa Cruz Biotechnology, sc-134220). After 24 h fresh virus was added. After 48 h the virus was removed, and the cells were allowed to recover overnight. Cell lines were selected using fluorescence-activated cell (FACs) sorting conducted by the MSK Flow Cytometry Core Facility. TIDE (Tracking of Indels by Decomposition) analysis was used to verify CRISPR-guided gene disruption^47^.

### Generation of stable cell lines

Genes encoding wild-type or mutant XPR1 were cloned into the lentiviral expression vector pLenti-GIII-CMV (Applied Biological Materials) and encodes the full-length protein followed by a C-terminal mCerulean tag. All constructs were validated by full plasmid sequencing. The cDNA expressing vectors along with lentiviral packaging plasmid psPAX2 and envelop plasmid pMD2.G-VSVG were transfected into LentiX cells using Lipofectamine 3000 transfection reagent (Thermo Scientific, L3000015). Virus-containing supernatant was collected 48 h after transfection and passed through a 0.45-μm filter. XPR1-knockout RPE-1 cells in 6-well tissue culture plates were infected with virus and 4 μg/ml Polybrene (Santa Cruz Biotechnology, sc-134220). After 24 h fresh virus was added. After 48 h the virus was removed, and the cells were allowed to recover overnight. Cell lines were selected by 20 μg/ml puromycin (Sigma, P8833). All cell lines were grown in DMEM medium (MSK Media Facility) containing 2 mM Pen/Strep supplemented with 10% fetal bovine serum (MSK Media Facility) and were maintained at 37°C and 5% CO_2_.

### Cloning, expression, and purification of XRBD

The XRBD of X-MLV envelope protein (strain NZB, Genebank #K02730, 32-238) fused to a C-terminal mouse Fc tag was synthesized by Twist Biosciences and cloned into a pcDNA vector with a N-terminal signal peptide (MKHLWFFLLLVAAPRWVLS). XRBD-mFc protein was expressed in Expi293F cells using transient transfection. After 4 days, the supernatant was collected and dialyzed into TBS buffer. Protein was purified using Protein G resin (GenScript, L00209), concentrated to 2.5 mg/ml in TBS buffer, and stored at −80°C.

### Radioactive Pi transport assays

Radioactive ^32^P substrate efflux experiments were carried out as follows. For FSEC analysis^48^, RPE-1 7.5 x 10^5^ cells were seeded per well in 2 ml DMEM medium in 6-well plates and for the transport assay, 2.5 x 10^5^ cells per well in 1 ml DMEM medium in 12-well plates. Both assays were performed 24 h later. The cells designated for analysis by FSEC were solubilized in 20 mM Tris, pH 7.5, 200 mM NaCl, and 1:1,000 dilution of Protease Inhibitor Cocktail Set III (EDTA free, Calbiochem) supplemented with 10 mM of the detergent LMNG (Anatrace). Samples were rotated for 1 h at 4°C and then centrifuged at 18,200 x g for 1 hr at 4°C before FSEC analysis. In parallel, to load the radioactive substrate, the cells designed for the transport assay were incubated in DMEM containing 0.5 µCi/mL [^32^Pi] (Revity, NEX053001MC) for 20 min at 37°C. Then, the cells were washed three times with DMEM medium. To initiate the efflux measurements, 400 µl of DMEM medium supplemented with 10 mM Pi, pH 7, was added. The reaction was carried out for 1 h at 37°C. An aliquot of medium was removed. Cells were washed three times in cold PBS (MSK Media Facility), and subsequently lysed using 1 ml of 1% Triton X-100 (Sigma, T9284). Two samples per well (250 µl of extracellular media; 750 µl of lysed cells) were transferred to a scintillation vial with 5 ml of scintillation cocktail (Perkin Elmer, 6013327). Radioactivity was measured with a liquid scintillation analyzer (Perkin Elmer, Tri-Carb 2910-TR). Efflux is graphed as a percentage of the [^32^Pi] that was accumulated by each cell type prior to the efflux assay using Graphpad Prism 10. All transport assay data are reported as normalized means ± standard deviation (with error propagation from the data transformation) from 3 replicates. The value for each cell line was normalized using the equation ((mutant-knock-out)/(wild-type/knock-out)). All mutants were tested in multiple, independent experiments.

### Cloning, expression, and purification of human XPR1

Human XPR1 was selected as a candidate for protein purification and structure determination using fluorescence-detection size-exclusion chromatography (FSEC) screening^48^. The full-length human XPR1 cDNA (synthesized by Twist Biosciences) was cloned into a modified pEG BacMam expression vector (addgene, 160686) containing a C-terminal mCerulean and Strep tag (WSHPNFEK) proceeded by a Precision Protease cleavage site (LEVLFQ/GP) to facilitate its removal.

XPR1 was expressed in Expi293F cells (Thermo Scientific A14527) using transient transfection. Briefly, 100 µg of plasmid DNA was incubated with 300 µg of polyethylenimine (PEI 25,000) (Polysciences, 23966-1) for 30 min at room temperature, then added to 100 ml of cells at a density of 2.0×10^6^ cells/ml. Cells were supplemented with 10 mM sodium butyrate after 20 h and harvested 44 h post-transfection and stored at −80°C.

A cell pellet was re-suspended in buffer containing 25 mM Tris, pH 7.5, 200 mM NaCl, a cOmplete Protease Inhibitor Cocktail Tablet (Roche). 1% lauryl maltose neopentyl glycol (LMNG, Anatrace, NG310) was added to the cell lysate and the mixture was rotated at 4°C for 1 h 30 min to extract XPR1 from membranes. The sample was centrifuged at 50,000g for 50 min at 4°C and the supernatant was filtered through a 0.22-μm polystyrene membrane (Millipore Sigma). The sample was incubated with Strep-Tactin XT resin (IBA Lifesciences) at 4°C for 1 h with rotation. At room temperature, beads were collected on a column, wash with buffering containing 25 mM Tris, pH 7.5, 200 mM NaCl, 0.002% LMNG (Anatrace, NG310) and 0.002% glycol-diosgenin (GDN, Anatrace, GDN101), and the protein as eluted with an identical buffer supplemented with 10 mM Biotin (IBA Lifesciences). The C-terminal mCerulean/Strep tag was removed by rotating the sample overnight at 4°C with PreScission Protease. XPR1 was further purified using a Superose 6 Increase size-exclusion column (Cytiva) in 25 mM Tris, pH 7.5, 200 mM NaCl, 0.002% LMNG and 0.002% GDN. For the InsP_6_-supplemented sample, 1 mM InsP_6_ (Sigma) was added to all purification buffers. For the high Pi- and InsP_6_-supplemented sample, all purification buffers contained 50 mM Na Phosphate, pH 7.4, and 150 mM NaCl (instead of 25 mM Tris, pH 7.5, and 200 mM NaCl) and 1 mM InsP_6_. For the InsP_8_-supplemented sample, 100 µM InsP_8_ (synthesized by the Jessen lab^28^) was added to the protein prior to gel filtration. For the Pi- and InsP_8_-supplemented sample, 5 mM Na Phosphate, pH 7.4, was added to all purification buffers and 100 µM InsP_8_ was added to the protein prior to gel filtration. Peak fractions were pooled and concentrated to 8-12 mg/ml using a 100-kDa concentrator (Amicon Ultra, Millipore Sigma).

### Cryo-EM sample preparation and data acquisition

Cryo-EM grids were prepared using a FEI Vitrobot Mark IV (Thermo Fisher Scientific) by applying 3 μl of purified XPR1 to a glow-discharged holy carbon gold QUANTIFOIL 1.2/1.3 grid (400 mesh, Electron Microscopy Sciences) and blotting for 4 to 5 s at 4°C and 100% humidity, then plunge freezing in liquid ethane. Cryo-EM datasets were collected on FEI Titan Krios microscopes (Thermo Fisher Scientific) operated at 300 kV housed in the MSK Richard Rifkind Center for Cryo-EM. Images were recorded in an automated fashion on a K3 Summit Direct Electron Detector (Gatan) in super-resolution counting mode with a super-resolution pixel size of 0.413 Å (physical pixel size of 0.826 Å) at a dose rate of 15 e^-^/pixel/s using SerialEM or a Falcon 4i Direct Electron Detector (Thermo Fisher) with a Selectris X Energy Filter (Thermo Scientific) in counting mode with a physical pixel size of 0.725 Å at a dose rate of 11.6 e^-^/pixel/s using EPU. The defocus range was −0.7 to −1.7 µm. Data collection statistics are shown in Table S1.

### Electron microscopy data processing

Image processing was performed using cryoSPARC (v4.5.0 or v4.5.1)^49^. The videos were gain-corrected, Fourier-cropped by two (0.826 Å) (when using the K3 camera only), and aligned using whole-frame and local-motion-correction algorithms within cryoSPARC. The cyroSPARC validation tool was used for FSC calculations and reliable resolution estimation.

#### Apo sample

Figure S1 shows the cryo-EM workflow for this dataset. A total of 5,500 super-resolution movies of apo XPR1 were collected using a K3 camera. Blob-based autopicking was implemented to select initial particles. Several rounds of two-dimensional (2D) classification were carried out and the best 2D classes (those with easy to identify transmembrane helices) were manually selected to generate two initial 3D models – the dimer and rotated dimer. False-positive selections and contaminants were excluded through iterative rounds of heterogeneous classification using the models generated from the ab initio algorithm as well as several decoy classes generated from noise particles through ab initio reconstruction. Particles contributing to these dimer and rotated dimer maps were combined and used for Topaz training and picking using neural networks, resulting in 4,266,672 particles^50^. Heterogeneous classification of these particles, followed by 3D classification using a focus mask covering the XPR1 density but excluding the detergent micelle density allowed separation of the best-aligning particles for both the dimer and rotated dimer. Stacks of 431,220 particles for the dimer and 124,969 particles for the rotated dimer were refined using non-uniform refinement and local refinement, yielding reconstructions of 2.52 Å and 3.06 Å, respectively. These final reconstructions were used as references for heterogeneous classification in future datasets.

#### InsP_6_-supplemented sample

Figure S2 shows the cryo-EM workflow for this dataset. A total of 4,116 super-resolution movies of InsP_6_-supplemented XPR1 were collected using a K3 camera. Blob-based autopicking was implemented to select initial particles. False-positive selections and contaminants were excluded through iterative rounds of heterogeneous classification using the models generated from apo sample as well as several decoy classes generated from noise particles through ab initio reconstruction. Particles contributing to this map were used for Topaz training and picking using neural networks, resulting in 5,084,405 particles^50^. Heterogeneous classification of these particles, followed by 3D classification using a focus mask covering the XPR1 density but excluding the detergent micelle density allowed separation of the best-aligning particles. A stack of 487,706 particles was refined using non-uniform refinement, yielding a reconstruction of 2.97 Å.

#### High Pi- and InsP_6_-supplemented sample

Figure S3 shows the cryo-EM workflow for this dataset. A total of 3,981 super-resolution movies of high Pi- and InsP_6_-supplemented XPR1 were collected using a K3 camera. Blob-based autopicking was implemented to select initial particles. Several rounds of two-dimensional (2D) classification were carried out and the best 2D classes (those with easy to identify SPX domains and transmembrane helices) were manually selected to generate initial 3D models. False-positive selections and contaminants were excluded through iterative rounds of heterogeneous classification using the models generated as well as several decoy classes generated from noise particles through ab initio reconstruction. Particles contributing to this map were used for Topaz training and picking using neural networks, resulting in 5,425,183 particles^50^. 2D classification and heterogeneous classification of these particles, followed by 3D classification using a focus mask covering the density for the SPX domain only allowed for separation of the best-aligning particles. A stack of 100,408 particles was refined using non-uniform refinement, yielding a reconstruction of 3.30 Å. This subset of particles gave a reconstruction with the SPX domain resolved, but at low resolution and in an orientation likely to be non-physiological.

#### InsP_8_-supplemented sample

Figure S4 shows the cryo-EM workflow for this dataset. A total of 12,900 movies of InsP_8_-supplemented XPR1 were collected using a Falcon 4i camera. Blob-based autopicking was implemented to select initial particles. Several rounds of two-dimensional (2D) classification were carried out and the best 2D classes (those with easy to identify SPX domains and transmembrane helices) were manually selected to generate an initial 3D model. False-positive selections and contaminants were excluded through iterative rounds of heterogeneous classification using the models generated from the ab initio algorithm as well as several decoy classes generated from noise particles through ab initio reconstruction. Particles contributing to this map were used for Topaz training and picking using neural networks, resulting in 5,312,652 particles^50^. Heterogeneous classification of these particles, followed by 3D classification using a focus mask covering the XPR1 density but excluding the detergent micelle density allowed separation of the best-aligning 934,621 particles. 3D classification, both with and without symmetry expansion, using a focus mask covering the XPR1 density but excluding the detergent micelle density allowed for separation of two unique C-terminal tail conformations – inward-open and occluded. Polished particle stacks were used for non-uniform refinement with global contrast transfer function (CTF) estimation, as well as higher-order tetrafoil, anisotropic magnification, and aberration corrections, local refinement, and reference based motion correction. The final reconstructions of the inward-open/occluded mixed dimer, 117,620 particles, and the occluded dimer, 295,448 particles, were 2.92 Å and 2.75 Å, respectively.

#### Pi- and InsP_8_-supplemented sample

Figure S5 shows the cryo-EM workflow for this dataset. A total of 13,132 movies of Pi- and InsP_8_-supplemented XPR1 were collected using a Falcon 4i camera. Data processing was analogous to the InsP_8_-supplement sample. The final reconstructions of the inward-open/occluded mixed dimer, 116,612 particles, and the occluded dimer, 338,543 particles, were 3.10 Å and 2.81 Å, respectively.

### Model building, refinement, and validation

ModelAngelo was used to automatically build initial atomic models into the cryo-EM density maps^51^. Subsequently, the models were manually rebuilt using Coot to fit the density^52^. Atomic coordinates were refined against the density-modified map using PHENIX real space refinement with geometric and Ramachandran restraints maintained throughout^53^. Model validation was performed using MolProbity^54^. Molecular graphics figures were prepared using UCSF Chimera^55^ or PyMOL. For electrostatic calculations, the APBS plugin in PyMOL was used^56^. To analyze and visualize tunnels in the XPR1 structures that lead from the Pi-binding sites to the surrounding solvent, MOLEonline was used^57^.

## Notes

### Competing Interest Statement

The authors have declared no competing interest.

