## Supplemental Document for "Transport and InsP8 activation mechanisms of the human inorganic phosphate exporter XPR1"

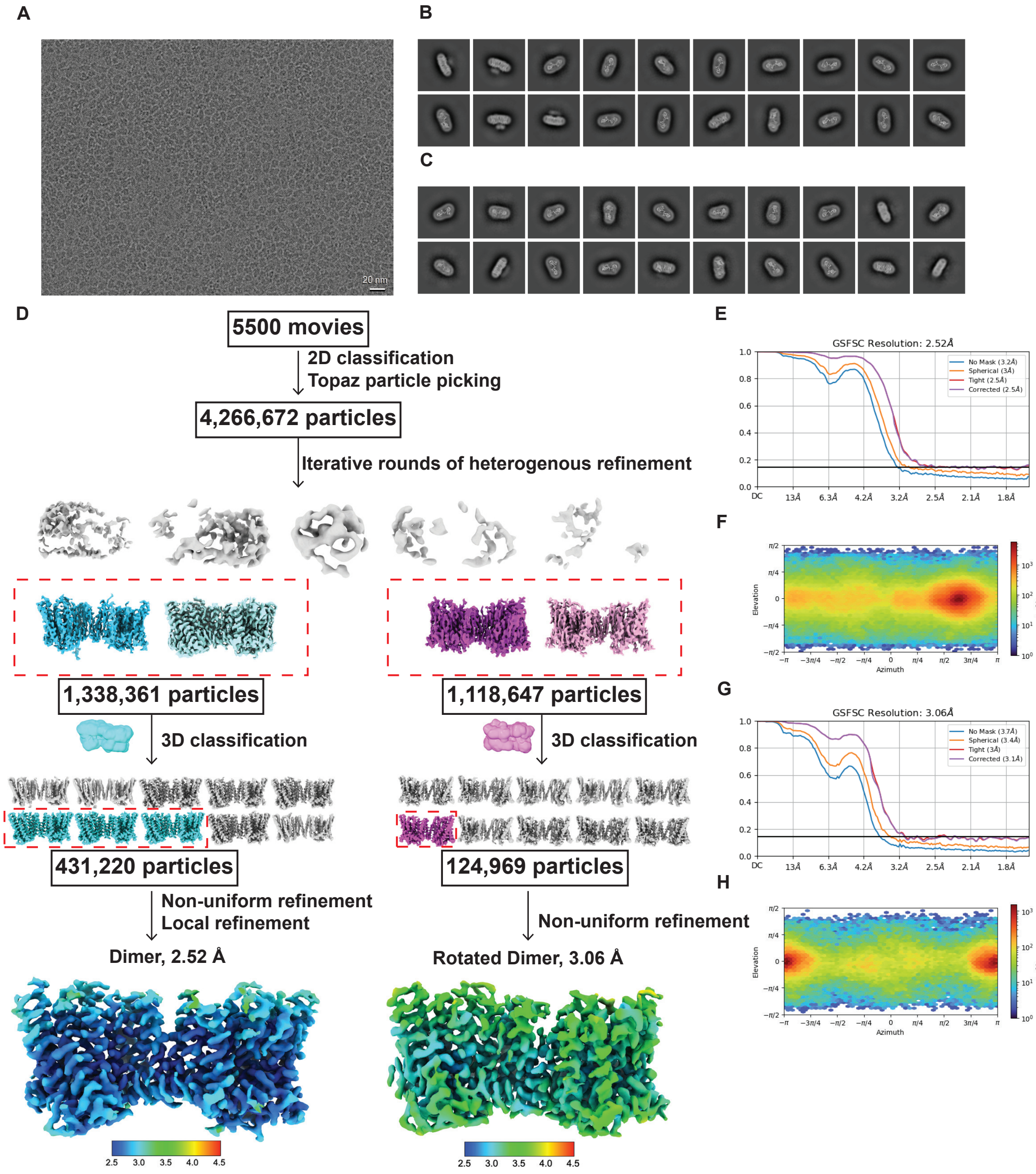

**Supplemental Figure 1. Cryo-EM data processing of apo XPR1. (A)** Representative micrograph. **(B)** Representative 2D class averages. **(C)** Representative 2D class averages of the rotated conformation. **(D)** Flowchart for cryo-EM data processing. **(E)** Gold-standard Fourier shell correlation (FSC) curve (cutoff of 0.143) of the final density map. **(F)** Angular orientation distribution of all particles used in the final 3D reconstruction. **(G)** Gold-standard FSC curve (cutoff of 0.143) of the final density map for the rotated conformation. **(H)** Angular orientation distribution of all particles used in the final 3D reconstruction for the rotated conformation.

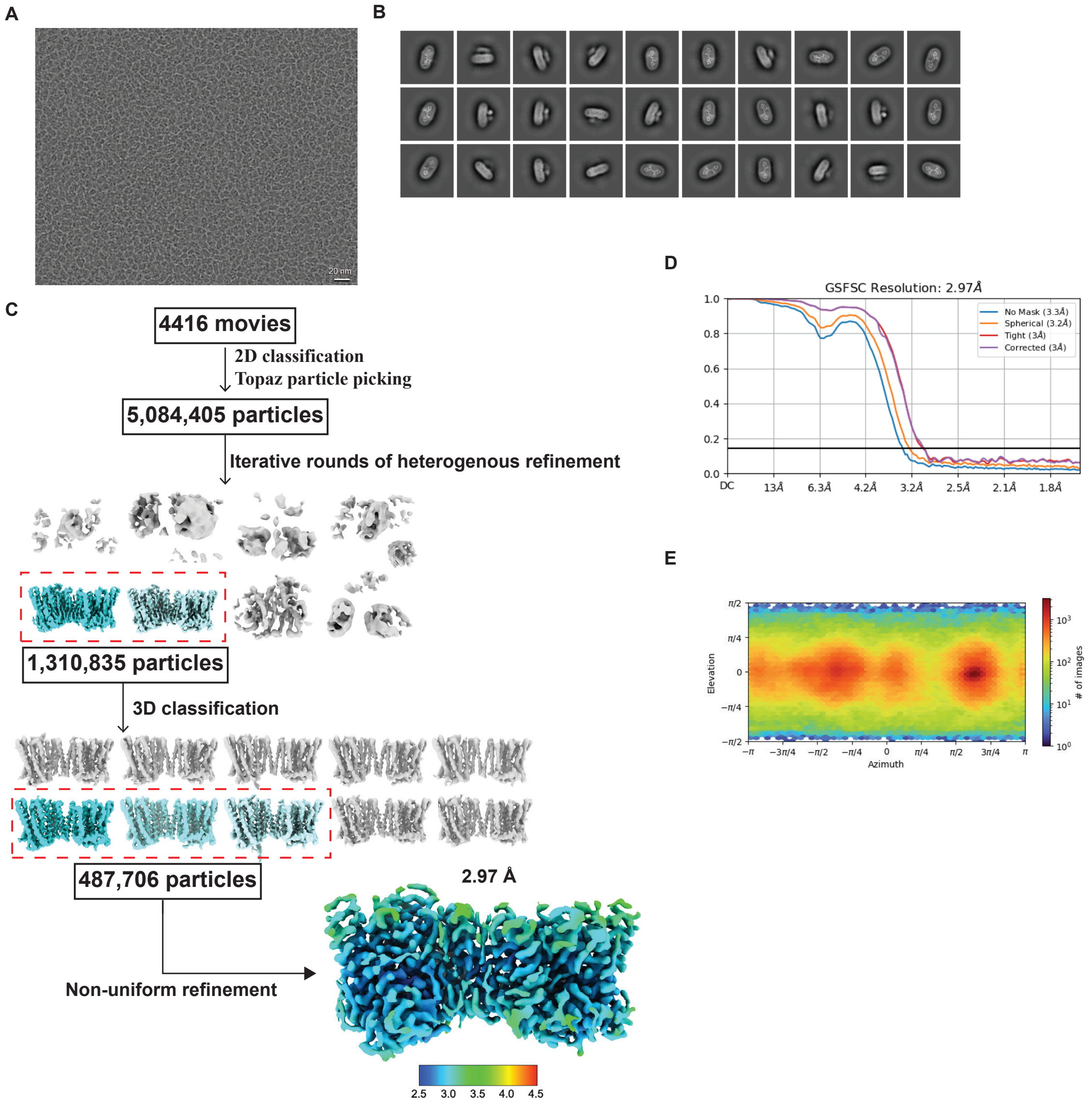

**Supplemental Figure 2. Cryo-EM data processing of $InsP_6$-supplemented XPR1. (A)** Representative micrograph. **(B)** Representative 2D class averages. **(C)** Flowchart for cryo-EM data processing. **(D)** Gold-standard FSC curve (cutoff of 0.143) of the final density map. **(E)** Angular orientation distribution of all particles used in the final 3D reconstruction.

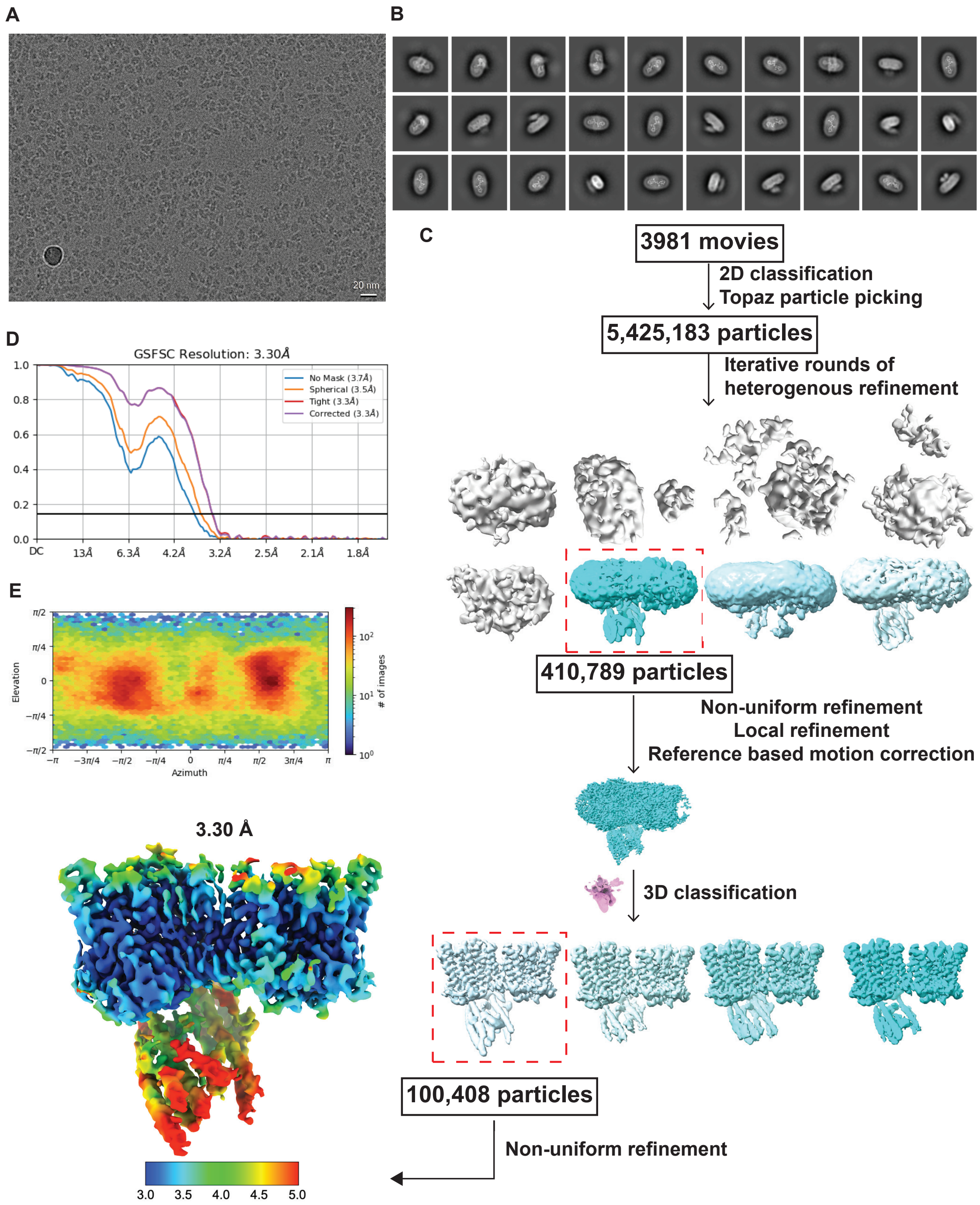

**Supplemental Figure 3. Cryo-EM data processing of high Pi- and InsP$_6$-supplemented XPR1. (A)** Representative micrograph. **(B)** Representative 2D class averages. **(C)** Flowchart for cryo-EM data processing. **(D)** Gold-standard FSC curve (cutoff of 0.143) of the final density map. **(E)** Angular orientation distribution of all particles used in the final 3D reconstruction.

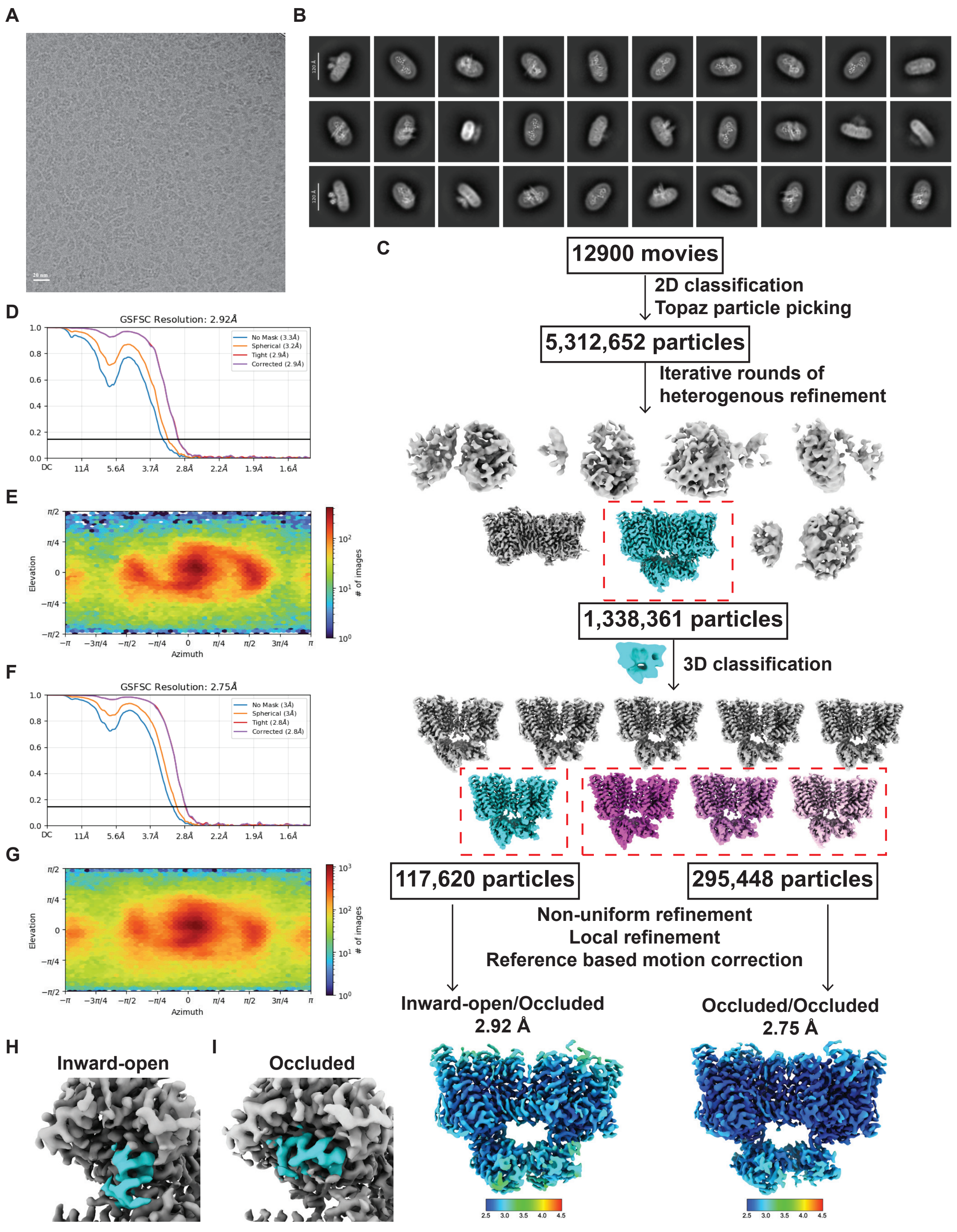

**Supplemental Figure 3. Cryo-EM data processing of InsP$_8$-supplemented XPR1. (A)** Representative micrograph. **(B)** Representative 2D class averages. **(C)** Flowchart for cryo-EM data processing. **(D)** Gold-standard FSC curve (cutoff of 0.143) of the final density map for the inward-open/occluded state. **(E)** Angular orientation distribution of all particles used in the final 3D reconstruction for the inward-open/occluded state. **(F)** Gold-standard FSC curve (cutoff of 0.143) of the final density map for the occluded state. **(G)** Angular orientation distribution of all particles used in the final 3D reconstruction for the occluded state. **H-I** C-terminal tail (cyan) position in the inward-open **(H)** and occluded **(I)** states.

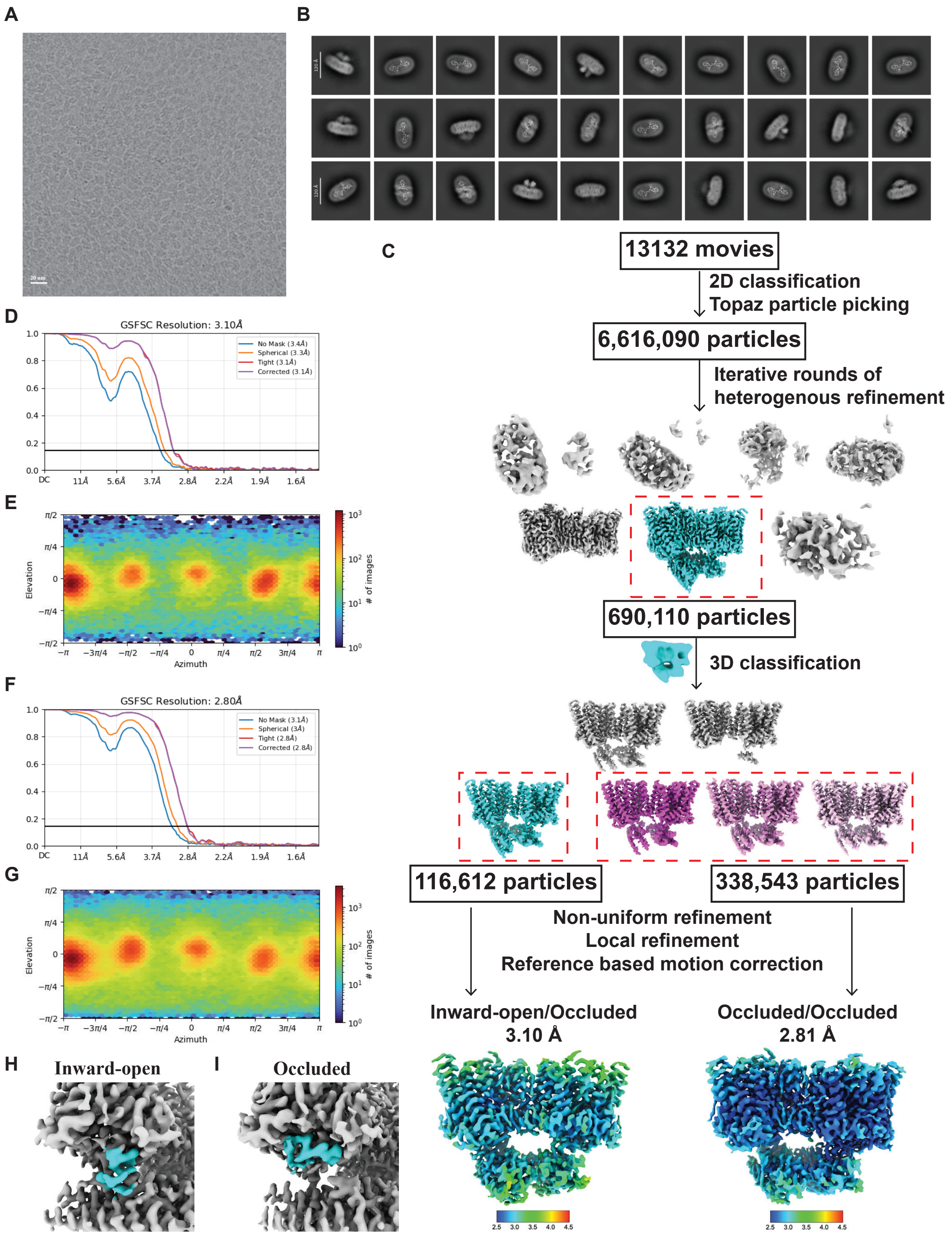

**Supplemental Figure 4. Cryo-EM data processing of Pi- and $InsP_8$-supplemented XPR1. (A)** Representative micrograph. **(B)** Representative 2D class averages. **(C)** Flowchart for cryo-EM data processing. **(D)** Gold-standard FSC curve (cutoff of 0.143) of the final density map for the inward-open/occluded state. **(E)** Angular orientation distribution of all particles used in the final 3D reconstruction for the inward-open/occluded state. **(F)** Gold-standard FSC curve (cutoff of 0.143) of the final density map for the occluded state. **(G)** Angular orientation distribution of all particles used in the final 3D reconstruction for the occluded state. **H-I** C-terminal tail (cyan) position in the inward-open **(H)** and occluded **(I)** states.

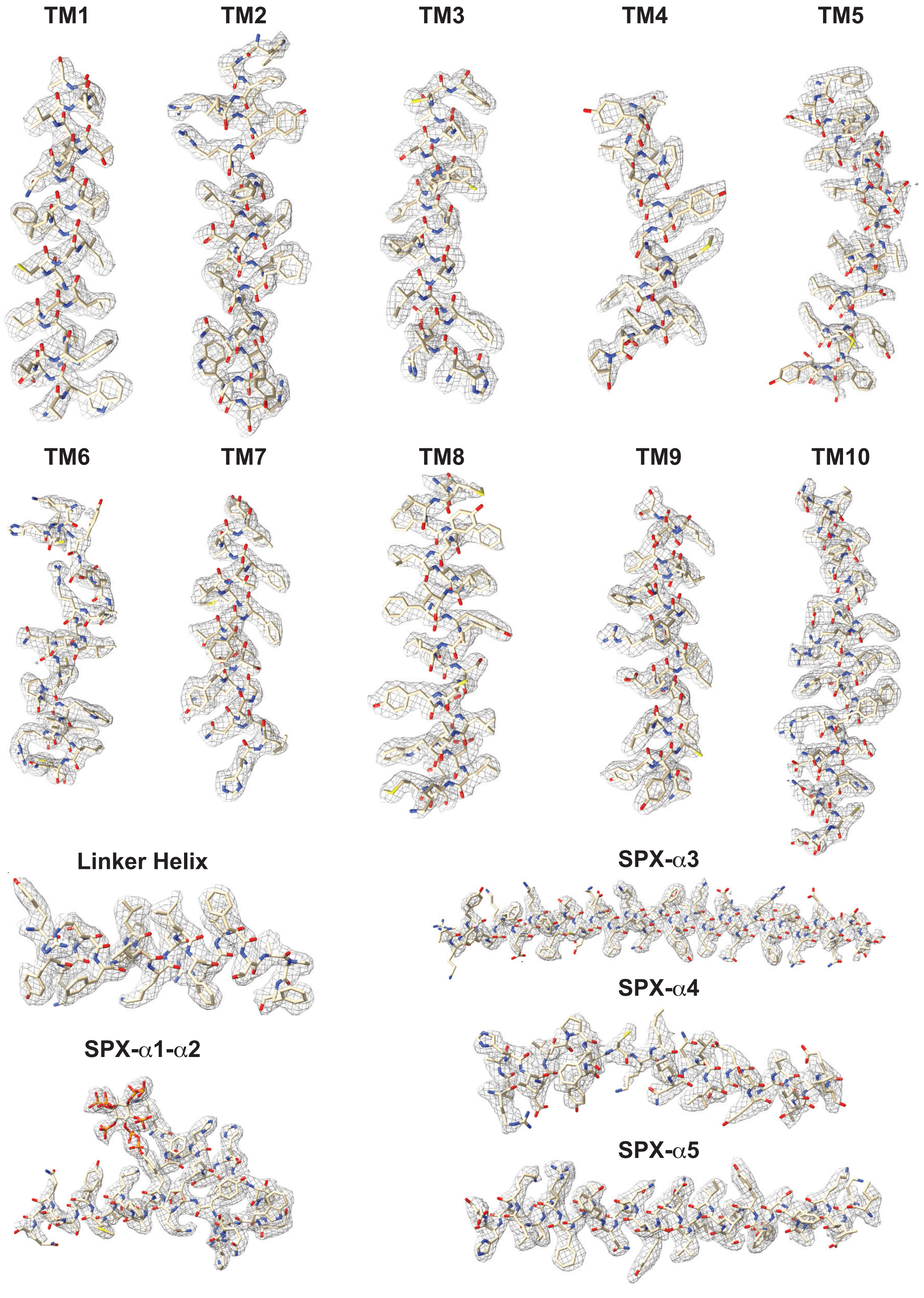

**Supplemental Figure 6. Quality of cryo-EM density of key structural elements.** Representative segments of the cryo-EM density map from the $InsP_8$-supplemented structure with the atomic models built *de novo*, with assistance from ModelAngelo. Each segment is labeled and demonstrates the quality of the map (contoured at the same level; gray mesh) from various regions of the reconstruction.

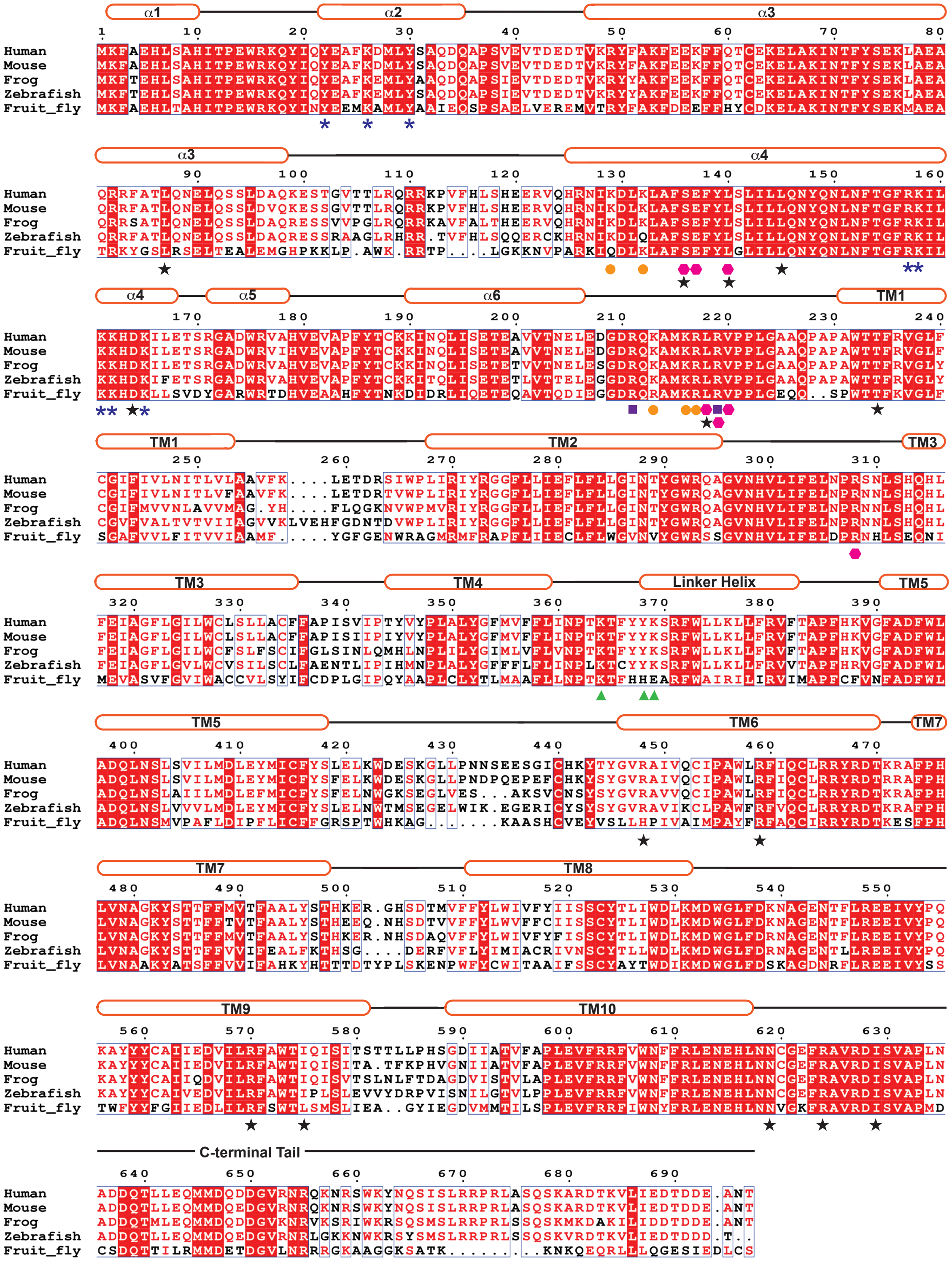

* : $InsP_8$ Site 1, SPX +ve Surface a
■ : $InsP_8$ Site 1, SPX +ve Surface b
● : $InsP_8$ Site 2, SPX +ve Surface c
▲ : $InsP_8$ Site 2, TM +ve Surface
⬢ : Arg Extension and Capture Pocket
★ : PFBC Missense Mutations

**Supplemental Figure 7. Sequence alignment of XPR1 orthologs.** The protein sequences of XPR1 from human (Uniprot - Q9UBH6), mouse (Uniprot - Q9QZ71), frog (Uniprot - Q28CY9), zebrafish (Uniprot - A8DZH4) and fruit fly (Uniprot - Q9VRR2) were aligned using the Clustal Omega server. The secondary structural is indicated with cylinders representing helices and solid lines representing loop regions. The alignment is colored according to conservation using the ESPript server. XPR1 residues identified in this study as important for $InsP_8$ binding and additional stabilization of the SPX domain are indicated: blue asterisks mark residues involved in $InsP_8$ binding site 1 on SPX +ve surface a, purple squares mark residues involved in $InsP_8$ binding site 1 on SPX +ve surface b, orange circles mark residues involved in $InsP_8$ binding site 2 on SPX +ve surface c, green triangles mark residues involved in $InsP_8$ binding site 2 on the TM +ve surface, and magenta hexagons mark residues involved in the Arg extension and capture pocket. Black stars mark known PFBC missense mutations.

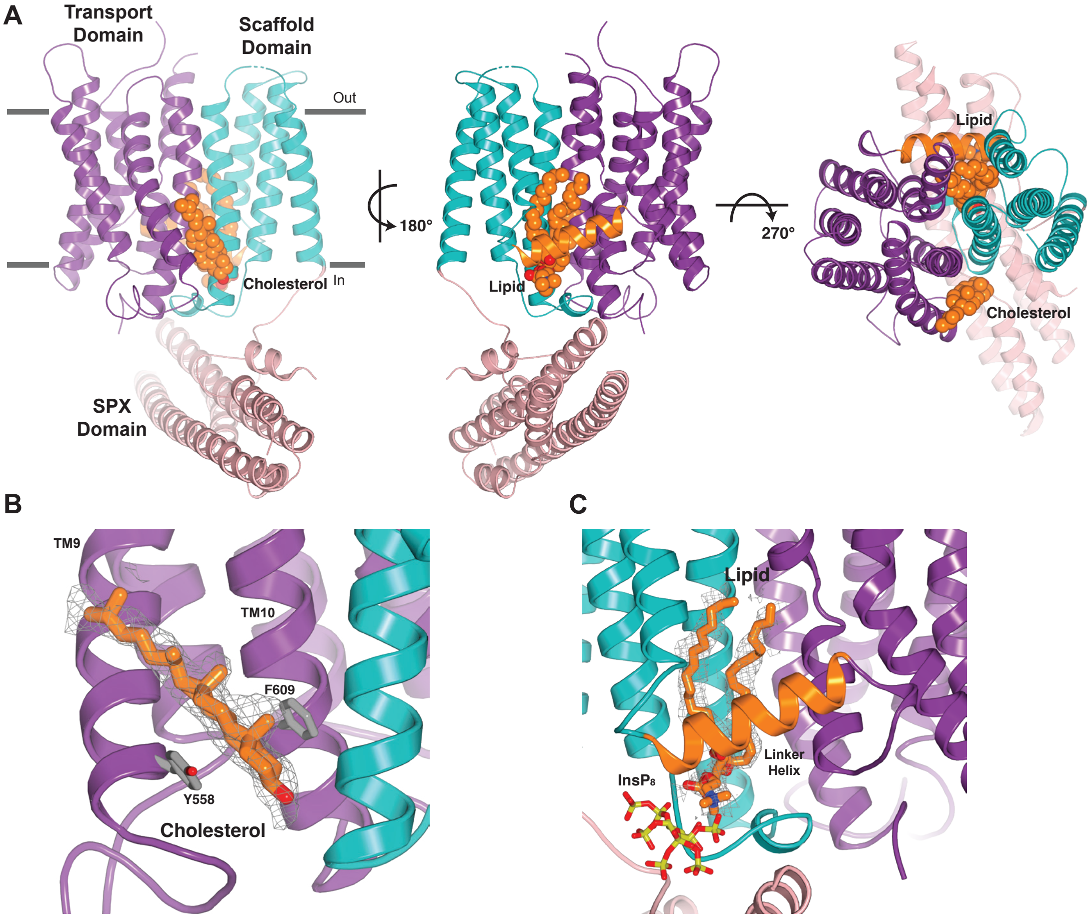

**Supplemental Figure 8. Lipid protein interactions between the scaffold and transport domains. (A)** Two well-ordered lipids, a cholesterol molecule and a phospholipid, pack between the scaffold and transport domains. The SPX domain is colored pink, the scaffold domain is colored teal, the linker domain is colored orange, and the transport domain is colored purple. **(B)** The cholesterol molecule is positioned by Tyr558 and Phe609. Cholesterol density is shown (gray mesh, 5σ contour). **(C)** The phospholipid was modeled as phosphatidyl choline, though its identity is not clear, and the density suggests that it could represent a mixture of lipid species. Phospholipid density is shown (gray mesh, 5σ contour).

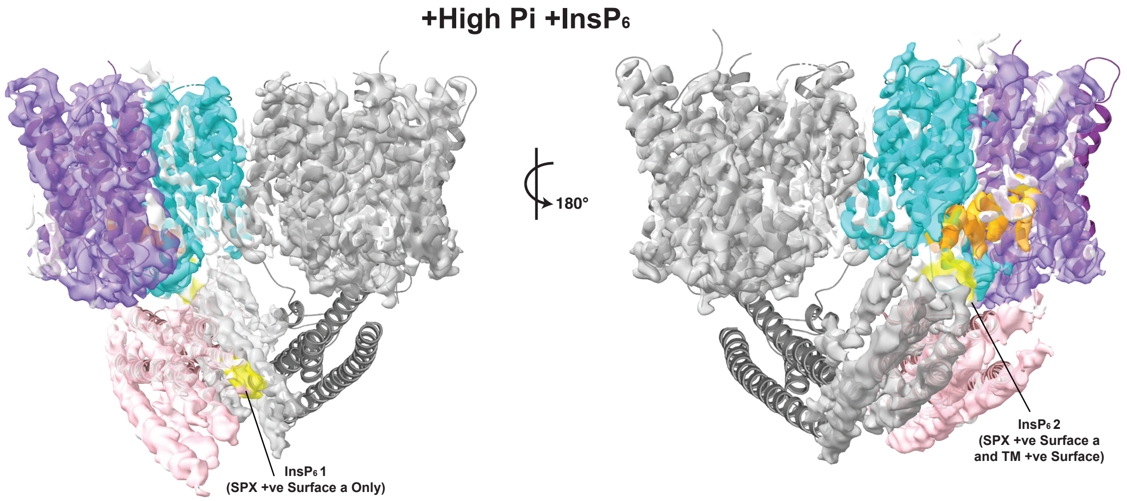

**Supplemental Figure 9. Structure of XPR1 in the presence of high Pi and $InsP_6$.** Cryo-EM density map of the XPR1 dimer, one monomer colored identically to Figure 1 and the other monomer colored grey, in the presence of high Pi and $InsP_6$. The bound $InsP_6$ is yellow. Overlay of the model of $InsP_8$-bound XPR1 illustrates the differences in SPX domain positioning.

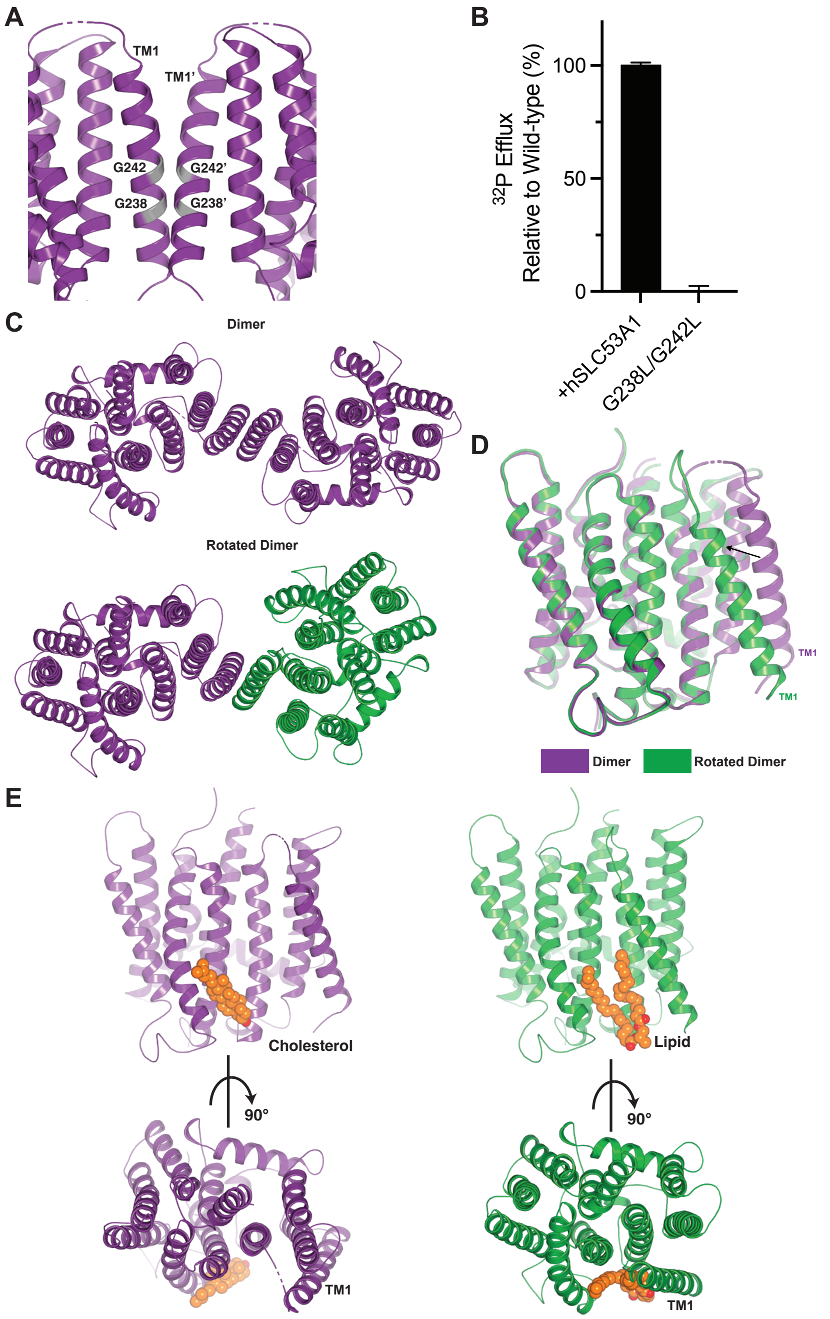

**Supplemental Figure 10. Dimerization of XPR1. (A)** The GxxxG packing motif at the dimer interface. **(B)** Export of radioactive $^{32}P$ by XPR1-knockout RPE-1 cells stably expressing wild-type or mutant XPR1-mCer where the GxxxG dimerization motif has been abolished. Data are represented as the normalized mean ± SD (n=3). **(C)** Two observed dimer configurations. **(D)** In the rotated dimer, TM1, which forms the dimer interface, has undergone a substantial movement in one of the two monomers leading to a rotation of monomer B with respect to monomer A. **(E)** The rearrangement of TM1 leads to the ejection of a well-defined cholesterol molecule.

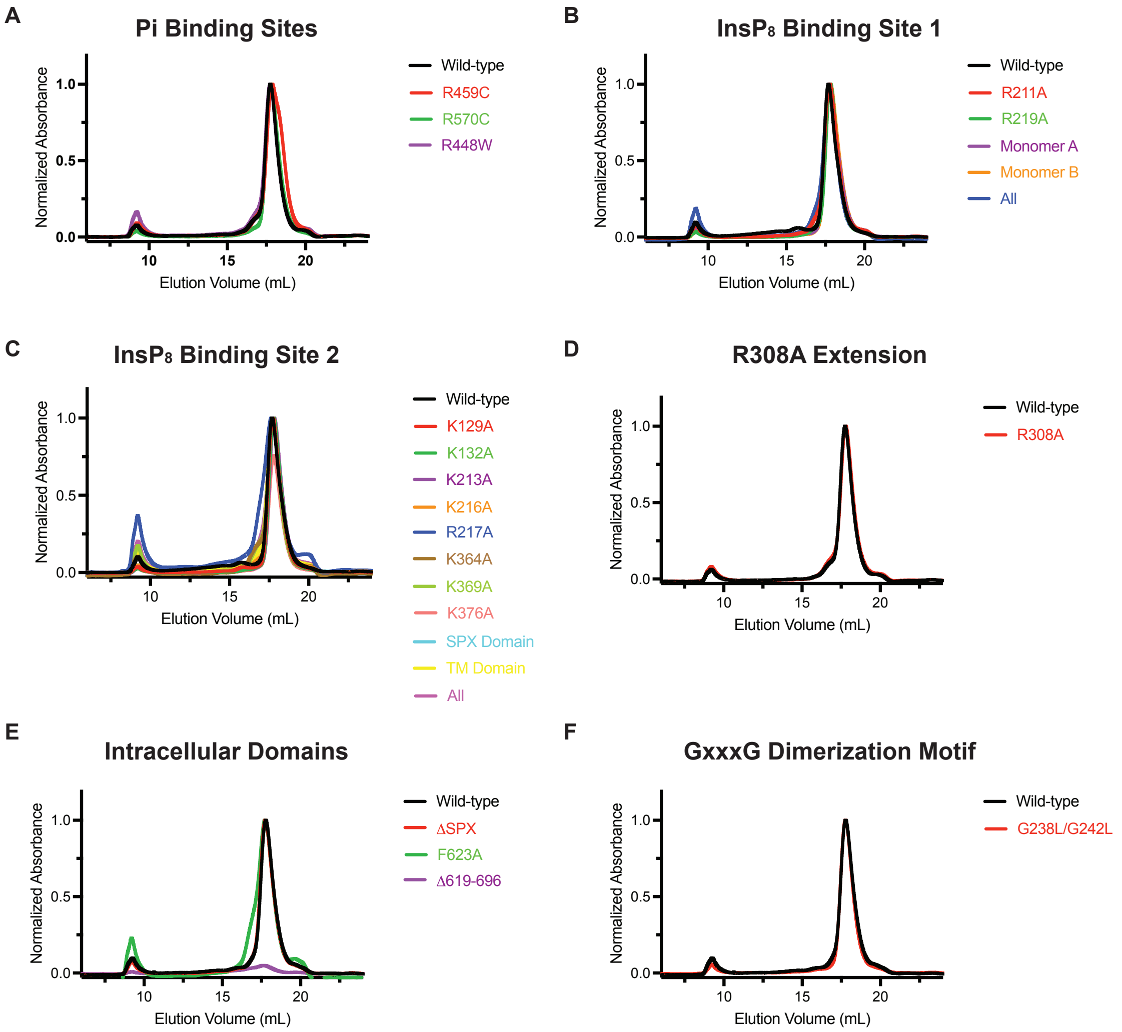

**Supplemental Figure 11. XPR1 wild-type and mutant behavior.** All XPR1 constructs were fused at the C-terminus to mCerulean and stably expressed in XPR1-knockout RPE-1 cells. **(A-F)** FSEC analysis of wild-type XPR1 and the mutants for which function was evaluated. Normalized FSEC traces are plotted for the Pi-binding site **(A)**, $InsP_8$-binding site 1 **(B)**, $InsP_8$-binding site 2 **(C)**, Arg extension **(D)**, intracellular domain **(E)**, and dimerization motif **(F)** mutants.

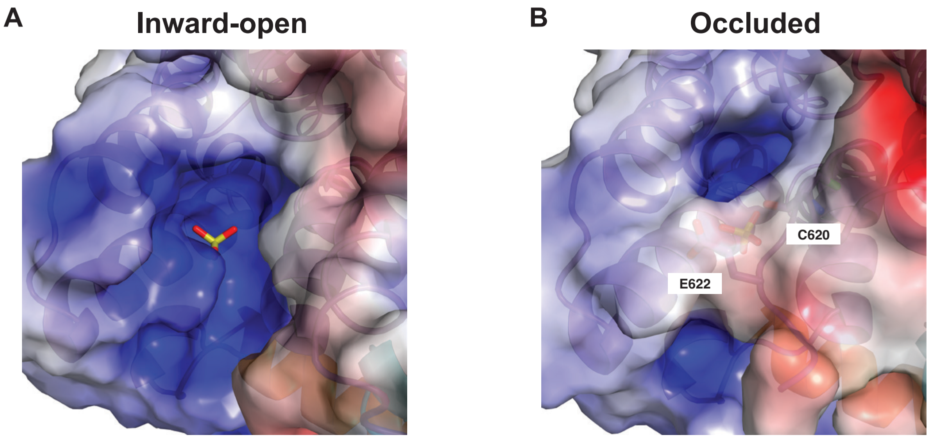

**Supplemental Figure 12. Electrostatics of the entrance to the substrate translocation pathway. (A)** In the inward-open state, the entrance is unobscured and positively charged. **(B)** In the occluded state, the C-terminal tail, and in particular Cys620 and Glu622, physically block the entrance to the substrate translocation pathway and introduce negative charge to repel the Pi substrate. The molecular surface is colored according to electrostatic potential: light gray regions are neutral; red, -5 $kTe^{-1}$; blue, +5 $kTe^{-1}$.

**Supplemental Data Table 1. Cryo-EM data collection, refinement, and validation statistics.**

| | #1 XPR1<br><br>(EMDB-xxxx)<br>(PDB xxxx) | #2 XPR1<br>Rotated Dimer<br>(EMDB-xxxx)<br>(PDB xxxx) | #3 XPR1<br>+InsP$_6$<br>(EMDB-xxxx)<br>(PDB xxxx) | #4 XPR1<br>+High Pi<br>+InsP$_6$<br>(EMDB-xxxx) |
|---|---|---|---|---|
| **Data collection and processing** | | | | |
| Magnification | 29,000 | 29,000 | 29,000 | 29,000 |
| Voltage (kV) | 300 | 300 | 300 | 300 |
| Electron exposure (e–/Å$^2$) | 66 | 66 | 66 | 66 |
| Defocus range (μm) | -0.7 to -1.7 | -0.7 to -1.7 | -0.7 to -1.7 | -0.7 to -1.7 |
| Pixel size (Å) | 0.826 | 0.826 | 0.826 | 0.826 |
| Symmetry imposed | C1 | C1 | C1 | C1 |
| Initial particle images (no.) | 4,266,672 | 4,266,672 | 5,084,405 | 5,425,183 |
| Final particle images (no.) | 431,220 | 124,969 | 487,706 | 100,408 |
| Map resolution (Å) | 2.52 | 3.06 | 2.97 | 3.30 |
| FSC threshold | 0.143 | 0.143 | 0.143 | 0.143 |
| Map resolution range (Å) | 2.35-8.07 | 2.7-13.0 | 2.55-8.40 | 2.79-9.54 |
| **Refinement** | | | | |
| Initial model used (PDB code) | ModelAngelo | ModelAngelo | ModelAngelo | |
| Model resolution (Å) | 3.00 | 3.21 | 2.98 | |
| FSC threshold 0.5 | | | | |
| Map sharpening *B* factor (Å$^2$) | -48.6 | -86.3 | -111.5 | |
| Model composition | | | | |
| Non-hydrogen atoms | 6470 | 6520 | 6472 | |
| Protein residues | 756 | 760 | 756 | |
| Ligands | 6 | 6 | 10 | |
| *B* factors (Å$^2$) | | | | |
| Protein | 95.68 | 84.98 | 56.70 | |
| Ligand | 93.68 | 87.24 | 54.73 | |
| R.m.s. deviations | | | | |
| Bond lengths (Å) | 0.006 | 0.007 | 0.005 | |
| Bond angles (°) | 0.691 | 0.769 | 0.645 | |
| Validation | | | | |
| MolProbity score | 2.01 | 2.21 | 2.08 | |
| Clashscore | 16.99 | 27.15 | 20.65 | |
| Poor rotamers (%) | 0.75 | 1.04 | 0.75 | |
| Ramachandran plot | | | | |
| Favored (%) | 95.97 | 96.00 | 96.10 | |
| Allowed (%) | 4.03 | 3.73 | 3.90 | |
| Disallowed (%) | 0.00 | 1.04 | 0.00 | |

| | #5 XPR1 +$InsP_8$ Inward-open/ Occluded (EMDB-xxxx) (PDB xxxx) | #6 XPR1 +$InsP_8$ Occluded/ Occluded (EMDB-xxxx) (PDB xxxx) | #7 XPR1 +Pi +$InsP_8$ Inward-open/ Occluded (EMDB-xxxx) (PDB xxxx) | #8 XPR1 +Pi +$InsP_8$ Occluded/ Occluded (EMDB-xxxx) (PDB xxxx) |
|---|---|---|---|---|
| **Data collection and processing** | | | | |
| Magnification | 165,000 | 165,000 | 165,000 | 165,000 |
| Voltage (kV) | 300 | 300 | 300 | 300 |
| Electron exposure (e–/$Å^2$) | 60.45 | 60.45 | 60.45 | 60.45 |
| Defocus range (μm) | -0.7 to -1.7 | -0.7 to -1.7 | -0.7 to -1.7 | -0.7 to -1.7 |
| Pixel size (Å) | 0.725 | 0.725 | 0.725 | 0.725 |
| Symmetry imposed | C1 | C1 | C1 | C1 |
| Initial particle images (no.) | 5,312,652 | 5,312,652 | 6,616,090 | 6,616,090 |
| Final particle images (no.) | 117,620 | 295,448 | 116,612 | 338,543 |
| Map resolution (Å) | 2.92 | 2.75 | 3.10 | 2.81 |
| FSC threshold | 0.143 | 0.143 | 0.143 | 0.143 |
| Map resolution range (Å) | 2.41-9.01 | 2.27-6.25 | 2.51-4.38 | 2.32-7.30 |
| **Refinement** | | | | |
| Initial model used (PDB code) | ModelAngelo | ModelAngelo | ModelAngelo | ModelAngelo |
| Model resolution (Å) | 2.86 | 2.64 | 2.97 | 2.71 |
| FSC threshold 0.5 | | | | |
| Map sharpening *B* factor ($Å^2$) | -77.9 | -88.2 | -83.6 | -87.6 |
| Model composition | | | | |
| Non-hydrogen atoms | 9962 | 9901 | 9950 | 9896 |
| Protein residues | 1153 | 1145 | 1153 | 1145 |
| Ligands | 16 | 16 | 14 | 15 |
| *B* factors ($Å^2$) | | | | |
| Protein | 16.07 | 7.11 | 56.70 | 7.31 |
| Ligand | 38.43 | 22.12 | 75.02 | 26.58 |
| R.m.s. deviations | | | | |
| Bond lengths (Å) | 0.005 | 0.004 | 0.004 | 0.004 |
| Bond angles (°) | 0.551 | 0.470 | 0.497 | 0.500 |
| Validation | | | | |
| MolProbity score | 1.95 | 1.97 | 2.05 | 2.05 |
| Clashscore | 23.72 | 20.87 | 27.97 | 25.44 |
| Poor rotamers (%) | 0.39 | 0.59 | 0.79 | 0.49 |
| Ramachandran plot | | | | |
| Favored (%) | 97.61 | 97.16 | 97.44 | 97.16 |
| Allowed (%) | 2.39 | 2.84 | 2.56 | 2.76 |
| Disallowed (%) | 0.18 | 0.00 | 0.00 | 0.09 |
